# OpenEMMU: a versatile, open-source EdU multiplexing methodology for studying DNA replication and cell cycle dynamics

**DOI:** 10.1101/2025.01.17.633495

**Authors:** Osvaldo Contreras, Chris Thekkedam, John Zaunders, Ismael Aguirre-MacLennan, Nicholas J. Murray, Anai Gonzalez-Cordero, Richard P. Harvey

## Abstract

5-Ethynyl-2’-deoxyuridine (EdU) has transformed DNA replication and cell cycle analyses through fast and efficient click chemistry detection. However, commercial EdU kits are expensive, contain proprietary reagents, are suboptimal for antibody multiplexing, and are challenging to use with larger biological specimens, highlighting the need for innovation. Here, we report Open-source EdU Multiplexing Methodology for Understanding DNA replication dynamics (OpenEMMU), an optimized, affordable, and user-friendly click chemistry resource using off-the-shelf reagents. OpenEMMU improves the efficiency, brightness, and multiplexing capabilities of EdU thymidine analog staining with both non-conjugated and conjugated antibodies across various cell types. We validated its effectiveness for fluorescent imaging of nascent DNA in developing embryos and organs, including the embryonic heart and forelimbs, and in 3D hiPSC-derived cardiac organoids. OpenEMMU also enabled 3D imaging of DNA synthesis in zebrafish larvae at fine resolution. This opens new avenues for insights into organismal development and growth, cell proliferation, and DNA replication, with precision and flexibility that surpass traditional methods.

**Teaser:** An open-source and cost-effective resource for DNA replication profiling and cell cycle analysis using click chemistry.

## INTRODUCTION

The cell cycle is a fundamental process in cell division, crucial for cell commitment, tissue morphogenesis, development, and homeostasis (Alberts et al., 2002). Understanding its core phases—G1, S, G2, and Mitosis—and their regulatory mechanisms is essential for understanding congenital malformations and adult diseases, including cancer (DePamphilis, 2006). Researchers use various tools and strategies to map the progression and phases of the cell cycle, including immunodetection of Ki67 for cell cycle competence (Sun & Kaufman, 2018), phosphorylated-Histone H3 for mitotic cells (Goto et al., 1999), Aurora B for cytokinesis (Goldenson & Crispino, 2015), nucleic acid dyes for DNA content analysis (Darzynkiewicz et al., 2010) and FUCCI reporters for cell cycle dynamics (Sakaue-Sawano et al., 2008).

DNA replication occurs during the S phase of the cell cycle, ensuring accurate duplication of genetic information (Sclafani, R. A. and Holzen, 2007). To study DNA replication, researchers have traditionally used thymidine analogs that are incorporated into newly synthesized DNA (Cavanagh et al., 2011). These analogs have provided insights into DNA replication rate, extent, and genome location, with thousands of studies demonstrating their use. Early methods used radioactive thymidine incorporation with autoradiographic detection (Ligasová and Koberna, 2018), whereas later, the incorporation of BrdU, a halogenated thymidine analog, was detected with antibodies (Gratzner, 1982; Nowakowski et al., 1989; Kuhn et al., 1996). More recently, EdU has gained popularity for its simpler, faster, and more sensitive detection using click chemistry (Fantoni et al., 2021; Ligasová and Koberna, 2018). Unlike BrdU detection, which uses harsh DNA denaturation conditions necessary for antigen retrieval, EdU click-chemistry-based detection minimizes cellular damage and other artifacts, is more efficient than traditional antibody-based approaches, and is well-suited for high-throughput studies, making EdU a preferred choice for DNA replication research (Darzynkiewicz et al., 2011; Ligasová and Koberna, 2018; Solius et al., 2021).

EdU detection utilizes click chemistry, a groundbreaking innovation that enables fast and efficient covalent conjugation of molecular entities (Fantoni et al., 2021). This technique was recognized with the 2022 Nobel Prize in Chemistry (Nobel Committee for Chemistry, 2022; Rostovtsev et al., 2002; Tornøe, Christensen, and Meldal, 2002). Specifically, the Cu(I)-Catalyzed Azide−Alkyne Cycloaddition (CuAAC) reaction demonstrates high catalytic efficacy when combined with reductants, proceeds rapidly and reliably under mild, often aqueous conditions (Rostovtsev et al., 2002; Fantoni et al., 2021), and achieves near-quantitative yields in forming stable triazole products (Meldal and Tornøe, 2008), making it ideal for biological systems. The specificity of the reaction is enhanced by the bioorthogonal azide and alkyne groups, and the stability of the triazole linkage, which is resistant to common chemical degradations, makes it suitable for long-term applications such as drug delivery and extended biomolecular labeling. Its versatility extends beyond biology, encompassing polymer science, drug and pharmaceutical development, and materials chemistry (Fantoni et al., 2021; Pang et al., 2022).

Since its introduction in 2008 (Salic & Mitchison, 2008), EdU has become a widely adopted tool for studying DNA replication, cell proliferation, and differentiation. When combined with DNA content assessment, it is particularly effective for identifying different cell cycle phases (Diermeier-Daucher et al., 2009). Notable examples of its application include studies on the proliferative nature of the stem cell gut niche (Carroll et al., 2018; Salic & Mitchison, 2008; Schepers et al., 2011), dividing stem cells (Solius et al., 2021), and adult skeletal muscle stem cell-mediated regeneration and quiescence (Cutler et al., 2022; Rocheteau et al., 2012). Additionally, EdU has been instrumental in research on embryonic and adult neurogenesis (Chehrehasa et al., 2009; Itaman, Enikolopov, & Podgorny, 2022; Lazutkin et al., 2019), the proliferative response of adult cardiac fibroblasts to myocardial damage (Soliman et al., 2020), and root plant growth (Hayashi, Hasegawa, & Matsunaga, 2013; Hsieh et al., 2015). EdU has also significantly contributed to cancer research and has been used to evaluate mammalian cardiomyocyte DNA synthesis and chromosome polyploidization after birth (Swift et al., 2023). Furthermore, it has been applied in super-resolution microscopy of DNA replication sites (Triemer et al., 2018; Li et al., 2021) and chromosome labeling (Ishizuka et al., 2016).

Recognizing its potential, several biotechnology companies have developed commercial EdU kits (e.g. Invitrogen by Life Technologies). However, EdU kits have several limitations that hinder scientific progress, including proprietary reagents and their concentrations, and high cost (USD 600-1,200 for 50 reactions/tubes). The need for separate kits for imaging and flow cytometry further limits flexibility and adds additional expenses when employing both techniques. Pipelines are not straightforward when using antibodies for parallel marker immunodetection, whether primary or conjugated, or other staining procedures and most EdU kits are limited to detecting 4-5 colors, restricting their multiplexing capabilities. In addition, methods are not optimized for use in whole organs or tissues in 3D, limiting analysis of larger and more complex biological structures or whole organisms. Overall, these limitations underscore the need for more affordable, open-source, and versatile alternatives, particularly for applications involving complex biological structures.

To address these limitations, we have developed <u>Open</u>-source <u>E</u>dU <u>M</u>ultiplexing <u>M</u>ethodology for <u>U</u>nderstanding DNA replication dynamics (OpenEMMU), a flexible and user-friendly clickable method, designed for high-parameter flow cytometry and various cutting-edge imaging modalities, utilizing commonly available and inexpensive reagents. We established OpenEMMU for fluorescent staining of clickable EdU in the adult gut stem cell niche, splenocytes, bone marrow cells, activated human T cells, whole developing organs including the embryonic heart and forelimb, and 3D self-assembled human organoids. Moreover, OpenEMMU was compatible with multiplexed imaging using “iterative bleaching extends multiplexity” or IBEX (Radtke et al., 2020) and enabled high-resolution 3D evaluation of nascent DNA synthesis in whole zebrafish larvae, which was not possible with traditional click staining methods. OpenEMMU allows for in-house optimization and validation at an unprecedented low cost, making it superior to commercially available EdU kits for DNA replication studies in simple and complex biological systems.

## RESULTS

### Optimizing OpenEMMU for S-Phase DNA replication profiling using CuAAC click chemistry

Since its discovery and application to the study of DNA synthesis and cell cycle dynamics (**Fig. 1A**) (Salic & Mitchison, 2008), EdU usage has steadily increased, while BrdU usage has substantially declined (**Fig. 1B**). As EdU detection relies on click chemistry, we aimed to develop an enhanced, open-source and cost-effective CuAAC-based EdU click reaction requiring common laboratory reagents and basic laboratory skills (**Fig. 1C**).

**Fig. 1.**
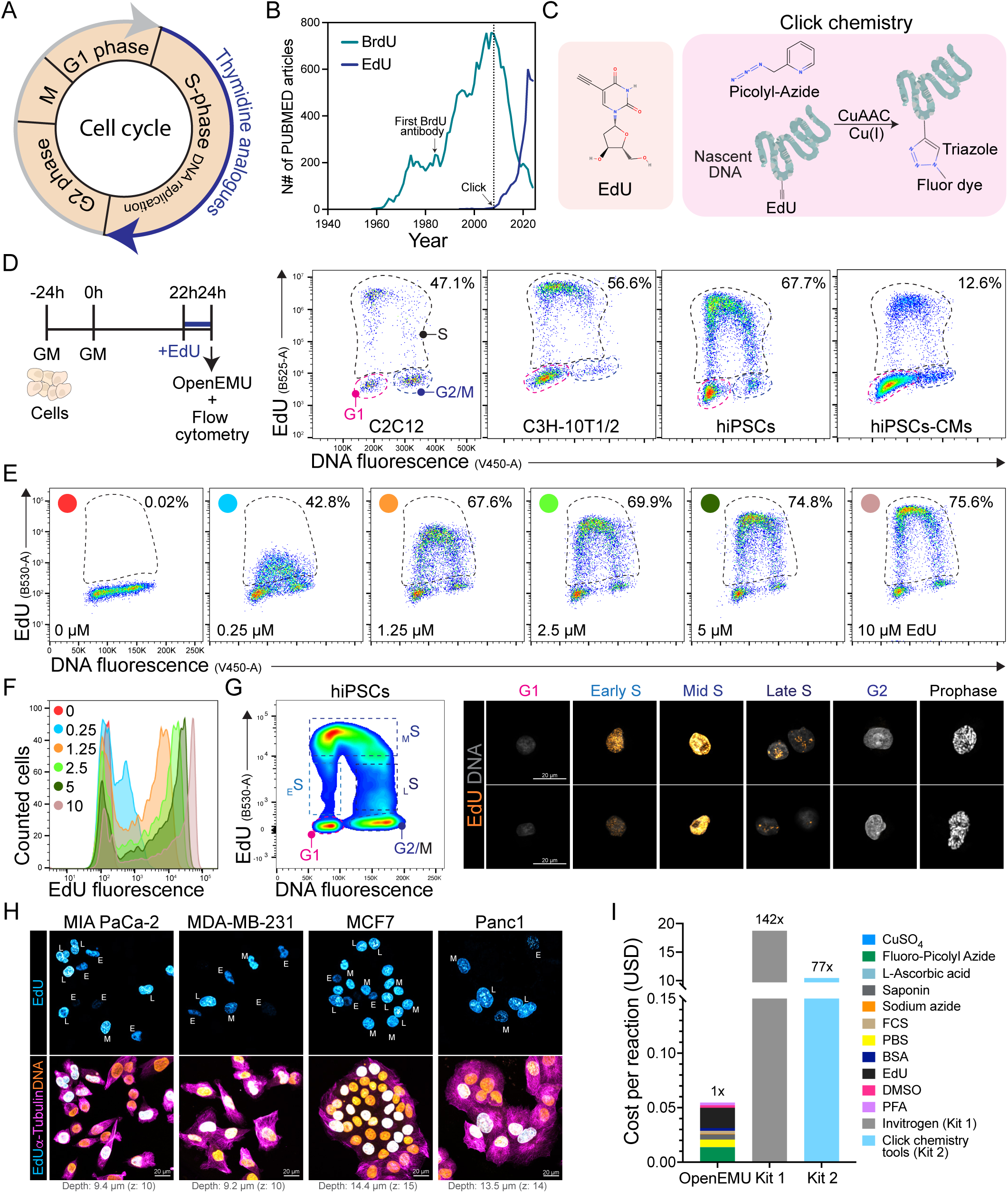
Application of OpenEMMU to normal and cancer cell lines for analysis of DNA replication. (A) Diagram outlining the cell cycle phases, highlighting the S phase where thymidine analogs are incorporated during DNA replication. (B) Graph depicting the annual number of PUBMED articles mentioning BrdU or EdU. (C) Illustration of the EdU molecule and CuAAC reaction in cells that have incorporated EdU into nascent DNA. (D) Flow cytometry results show the proportion of EdU-labeled cells after a 2-hour pulse in indicated cell lines (*n* = 3). (E) Relationship between EdU fluorescence intensity and the proportion of EdU+ cells at varying EdU concentrations in hiPSCs (*n* = 3). (F) Comparison of EdU and DNA fluorescence intensity at different EdU concentrations from (E). (G) Flow cytometry analysis of hiPSCs following a 1-hour EdU pulse. The dotted lines indicate the gating strategy employed for both FACS and subsequent confocal microscopy of sorted cells (right panels) (*n* = 3). (H) Confocal microscopy images of indicated EdU+ cancer cell lines after a 2-hour EdU pulse, with α-Tubulin immunolabeling and DNA staining (*n* = 3). E: Early DNA replication (S-phase); M: Mid DNA replication; L: Late DNA replication. (I) Cost per 1mL reaction comparing OpenEMMU against two commercially available kit options (Invitrogen, kit 1; Click Chemistry Tools, kit 2).

We conducted a systematic evaluation of critical reagents and reaction conditions, to refine CuAAC click chemistry for precise and efficient detection of nuclear DNA synthesis in proliferating cells. This analysis assessed their effects on labeling efficiency and signal quality across diverse cell lines and species (**Fig. 1 and Fig. S1**). To improve usability, we created detailed step-by-step guides for flow cytometry and imaging (**Materials and Methods; Fig. S2 and S3A**), incorporating antibody immunolabeling and immunohistochemistry. In our flow cytometry and imaging studies, cells were treated with EdU for 2h at increasing concentrations (0-10µM) and after fixation in PFA were incubated with an optimized OpenEMMU click reaction mixture, consisting of Cu-chelating azide dye (AZDye488/555/633/680 Picolyl Azide; 0.2 µM), copper catalyst (CuSO□.5H2O; 0.8 mM), and sodium ascorbate reducing agent (L-ascorbic acid; 1 mg/mL) in PBS for 30 minutes (**Materials and Methods; Fig. 1D and E**). Our evaluation showed that CuSO was a limiting reagent, and DNA replicating cells were not labeled properly below 800 µM (**Fig. 1E and F, Fig. S1A-C**). Whereas 400 µM produced a significant number of EdU-labeled cells, labeling efficiency was reduced compared to 800 µM in our standardized conditions (**Fig. S1C**). Increasing the CuSO concentration to 2 mM did not improve labeling but diminished DNA content fluorescence intensity as measured by Vybrant Violet™ DNA dye fluorescence (**Fig. S1C**).

To further optimize the CuAAC-based click reaction, we quantified the click reaction efficiency using L-ascorbic acid as the reducing agent. Among the tested concentrations, 0.5 mg/mL and higher were effective, leading us to select 1 mg/mL as our working concentration (**Fig. S1D**). 0.1 mg/mL did not facilitate the CuAAC click reaction. These findings highlight the importance of reducing agent concentration for efficient click chemistry (Meldal and Tornøe, 2008). We also investigated the efficiency of the copper-chelating organic picolyl azide in the CuAAC click reaction. Picolyl azides were effective at concentrations between 0.1–0.5 µM but caused overstaining of non-EdU cells at higher concentrations (2–10 µM) (**Fig. S1E**). Thus, increasing the concentration of Alexa Fluor-conjugated picolyl azide did not improve staining or separation but reduced the signal-to-noise ratio. Therefore, we selected 0.2 µM as the optimal working concentration for picolyl azides (tested AF488/555/633/680).

We then compared two bovine serum types in our adapted permeabilization and wash buffer, to further refine our protocol. Both Newborn Calf Serum (NCS) at 2% and Fetal Bovine Serum (FBS) at 4% effectively supported EdU labeling (**Fig. S1F**). NCS was selected due to its lower cost compared to FBS, although either can be used depending on researchers’ needs. We also explored various DNA dyes and observed that OpenEMMU was compatible with all tested. However, Vybrant™ DyeCycle™ Violet was significantly brighter than Hoechst 33342, Hoechst 33258, and DAPI, even at lower concentrations (5 µM Vybrant™ DyeCycle™ Violet versus 18 µM for the other DNA dyes) (**Fig. S1G**).

### OpenEMMU enables the cost-effective investigation of DNA replication in multiple cell types using flow cytometry and fluorescence microscopy

Two hours of EdU treatment at 10 µM effectively labels various cell types, including C2C12 myoblasts, C3H10T1/2 mesenchymal stromal cells, human-induced pluripotent stem cells (hiPSCs), and hiPSC-derived cardiomyocytes at day 12 of differentiation (**Fig. 1D, Fig. S3B**). These cell types exhibit varying degrees of DNA replication rate and S-phase cell proportions (approximately 47, 57, 68, and ∼13% of total cells, respectively). The horseshoe-shaped EdU/DNA bivariate distribution allows for the identification of cells in the G1 phase (2n DNA, no EdU), S phase (DNA replicating, EdU+), and the G2/M phases (4n DNA, no EdU) of the cell cycle by combining EdU incorporation and DNA fluorescence detection using Vybrant Violet (**Fig. 1D**). We confirmed that the optimal EdU concentration range of 5-10 µM is necessary for accurately labeling DNA-replicating hiPSCs, as lower concentrations (below 5 µM) significantly reduce EdU fluorescence intensity and the proportion of S-phase cells detected (**Fig. 1E and F**). Therefore, our unified clickable method delivers excellent results for both flow cytometry and fluorescence imaging, overcoming the limitations of most commercial EdU kits, which typically support only one of these techniques.

OpenEMMU also enabled fluorescence-activated cell sorting (FACS) of distinct DNA replicating cells, specifically early, mid, and late S-phase cells, using the horseshoe-shaped EdU/DNA bivariate fluorescence as a proxy (**Fig. 1G**). FACS-assisted separation followed by cytospin of EdU+ hiPSCs (1h EdU) allowed us to distinguish distinct DNA replicating hiPSC populations and identify specific regions of DNA synthesis within the nuclear DNA. Notably, the intensity of EdU fluorescence and DNA content allowed differentiation of compartmentalized DNA replication dynamics among early, mid, and late replicating cells (**Fig. 1G**). Open, less condensed euchromatic DNA replicates first, followed by more condensed heterochromatic DNA, which typically replicates during the late S-phase (Chagin, Stear & Cardoso; Fu, Baris & Aladjem, 2018). Early replicating DNA in hiPSCs undergoing the S-phase exhibited a less intense, dispersed, and punctate DNA synthesis pattern largely excluding the nucleolus, whereas mid-replicating cells displayed larger and more abundant DNA replication foci concentrated at the nuclear periphery and within the nucleoplasm. In contrast, late S-phase cells showed a patchier and more segregated DNA replication pattern, particularly within the nucleolus and other late-replicating regions and chromosomes (**Fig. 1G**). Co-staining with NUCLEOLIN (for nucleoli) and EdU in hiPSCs, using a brief 10-minute labeling period, enabled precise identification of replicative events at different stages of the S-phase (**Fig. S4A**). Thus, OpenEMMU significantly enhances S-phase cell profiling by allowing for high-resolution discrimination and evaluation of DNA synthetic spatial domains.

To explore other relevant cell types, we examined various cancer cell lines (Panc1, MIA PaCa-2, MCF7, and MDA-MB-231) and co-stained them with cytoskeletal α-Tubulin for confocal imaging (**Fig. 1H**). In these cells, OpenEMMU enabled the evaluation of DNA replication and the simultaneous flow cytometric analysis of EdU with mitotic Histone H3 phosphorylated at serine 10 (p-HH3^S10^) and DNA damage (γH2A.X^S139^) markers (**Fig. S4B-D**). Moreover, OpenEMMU effectively distinguished early, mid, and late S-phase cells based on the label intensity and characteristic genomic locations of DNA replication foci in cancer cell lines and hiPSCs (**Fig. 1H and Fig. S4**). We successfully co-stained 2-hour EdU-labeled hiPSCs with the pluripotency transcription factor OCT3/4, reinforcing OpenEMMU’s versatility for co-immunolabeling (**Fig. S4E**). To validate OpenEMMU’s accuracy in detecting DNA replication dynamically and measuring cell cycle progression, we stimulated DNA synthesis in C2C12 myoblasts and C3H10T1/2 mesenchymal stromal cells using FGF-2 and PDGF-BB ligands. Both growth factors induced DNA replication 3-4 times and promoted cell cycle progression in low-serum conditions, as measured by EdU incorporation and the horseshoe-shaped EdU/DNA bivariate fluorescence (**Fig. S4F**). While these cell lines demonstrated increased cell cycle progression in response to both growth factors, C2C12 myoblasts exhibited more robust proliferation with FGF-2, whereas C3H10T1/2 cells responded more favorably to PDGF-BB (**Fig. S4F**). This differential response is likely attributable to the distinct relative expression of PDGF receptor proteins (PDGFRα, PDGFRβ) in these cell types (Contreras et al., 2021), highlighting the ability of OpenEMMU to provide nuanced data on cell signaling-driven cell cycle responses.

For our optimized OpenEMMU method, we calculated the cost of each reagent, including those for fixation, permeabilization, and click chemistry of EdU-labeled cells, per milliliter of reaction. At the time of writing, with an estimated cost of < USD 0.05 per 1 mL of click chemistry reaction, OpenEMMU is approximately 142 and 77 times less expensive than the Invitrogen EdU Flow Cytometry and VectorLabs Click Chemistry Tools kits, respectively (**Fig. 1I**). Thus, our analysis shows that OpenEMMU can be assembled in-house cost-effectively using common, off-the-shelf reactants and reagents.

### OpenEMMU demonstrates superior detection and cell cycle analysis compared to commercial EdU kits across diverse cell models

We then compared directly the Invitrogen EdU kit for flow cytometry to OpenEMMU to quantitatively measure EdU fluorescence intensity and evaluate the accuracy of EdU fluorescence detection in S-phase cells. OpenEMMU consistently outperformed the Invitrogen kit in both aspects, accurately detecting a higher proportion of S-phase cells (by ∼5-10%, *n*=3) and exhibiting ∼10-fold greater EdU fluorescence intensity (**Fig. 2A and B, Fig. S5A**). This superiority was evident across a wide range of cell types, including highly proliferative undifferentiated hiPSCs and slowly DNA-replicating cells such as hiPSC-derived cardiomyocytes at day 20 of differentiation and the C2C12 myoblast cell line (**Fig. 2A and B, Fig. S5A**).

**Fig. 2.**
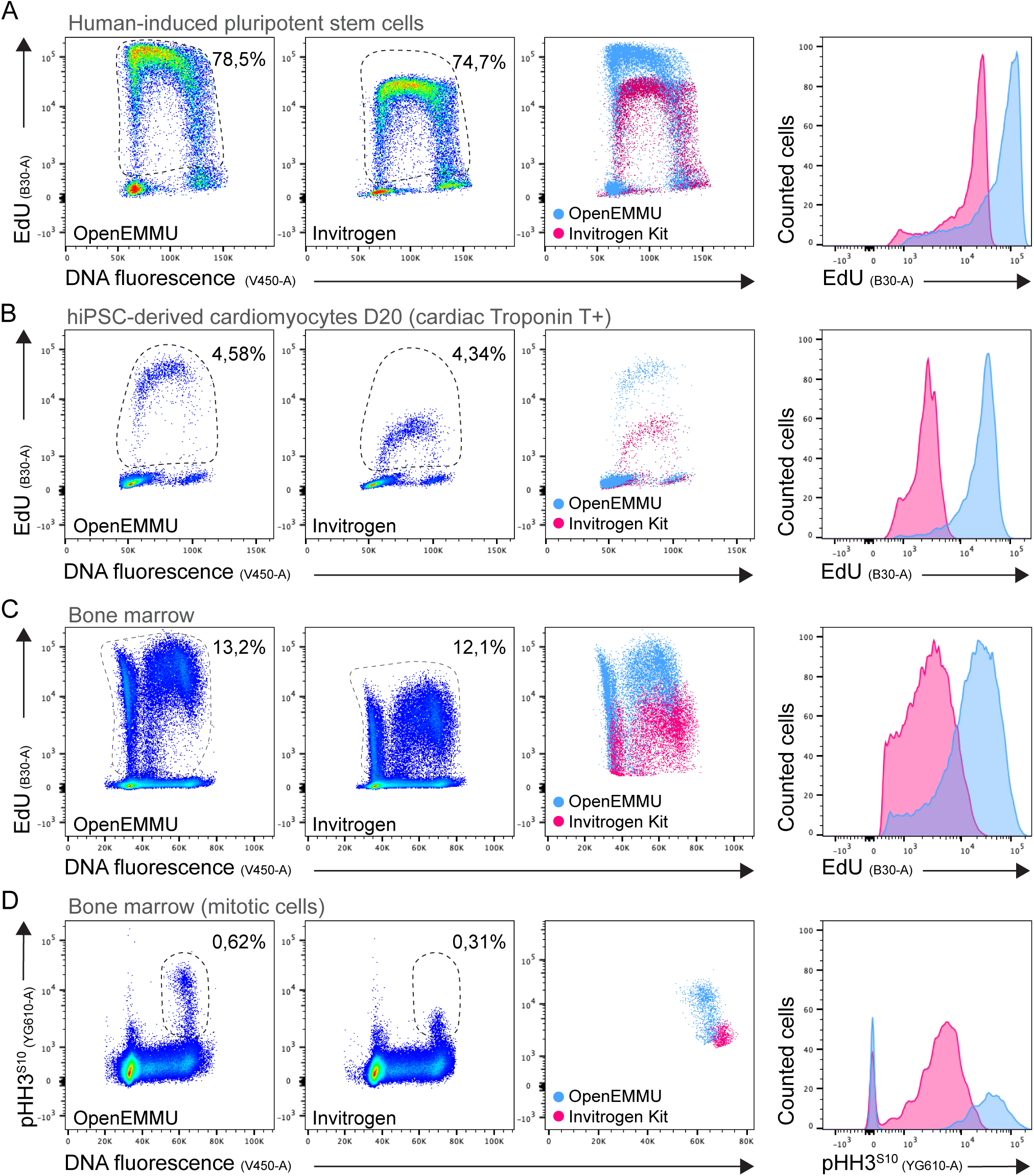
OpenEMMU outperforms commercial kits in evaluating DNA-replicating and mitotic cells. (A) Flow cytometry results comparing OpenEMMU and the Invitrogen kit in terms of EdU fluorescence intensity and the proportion of EdU-labeled cells after a 2-hour pulse in hiPSCs (*n* = 3). (B) Flow cytometry results comparing EdU fluorescence intensity and the proportion of EdU+ cells in hiPSC-derived cardiomyocytes (cTnT+) at day 20 of differentiation (*n* = 3). (C) Flow cytometry results comparing EdU fluorescence intensity and the proportion of EdU+ cells in total bone marrow cells of mice injected with EdU for 3 hours (*n* = 3). (D) Flow cytometry analysis of phosphorylated histone H3 at serine 10 (p-HH3^S10+^) to detect mitotic cells after click chemistry using OpenEMMU versus the Invitrogen EdU kit, demonstrating a significant reduction in detected mitotic cells with the Invitrogen kit (*n* = 3).

Beyond cell lines, we compared the performance of OpenEMMU and Invitrogen EdU kits using tissue harvested after *in vivo* EdU uptake, including bone marrow cells and splenocytes from healthy wild-type mice. Adult WT male and female mice were intraperitoneally injected with EdU for 4 hours (based on its bioavailability of approximately 1-2 hours (Maltsev et al., 2022)), followed by OpenEMMU analysis of PFA-fixed cells. In bone marrow and spleen cells, OpenEMMU accurately stained DNA-replicating cells and produced significantly brighter EdU-labeled cells (i.e., fluorescence intensity) (**Fig. 2C and Fig. S5B**). We then evaluated OpenEMMU’s compatibility with co-staining using a phycoerythrin (PE)-conjugated antibody to detect mitotic cells (p-HH3^S10+^). PE, a red protein pigment, is known to be adversely affected by CuAAC click chemistry in most commercial EdU kits (Glazer, 1990), and as expected, detection of mitotic cells using a PE-conjugated antibody using the Invitrogen kit was suboptimal (**Fig. 2D**). In contrast, OpenEMMU successfully labeled these cells, with a detection rate of 0.62% of mitotic cells compared to 0.31% using the Invitrogen EdU kit. Not only did the commercially available kit impact the proportion of p-HH3^S10+^ mitotic cells but also the fluorescence intensity of the PE-conjugated antibody (**Fig. 2D**). These results underscore OpenEMMU’s versatility across various cell models and staining procedures, making it a simple and adaptable EdU multiplexing methodology for studying DNA replication and cell cycle.

### OpenEMMU enables multiplexing with conjugated antibodies for immune analysis

To challenge OpenEMMU’s multiplexing capabilities further, we first combined the detection of DNA-replicating bone marrow and spleen adult cells using EdU with immune markers such as CD45, CD11c, and CD11b, analyzed via flow cytometry (**Fig. 3A-C, Fig. S6A**). OpenEMMU effectively detected not only highly proliferative bone marrow cells but also splenocytes (CD11b/c+), known for their much lower proliferative rate—these were found to represent 1% of proliferative cells within the total splenocyte population (**Fig. 3B and C**). Notably, OpenEMMU identified myeloid CD11b+ cells with an exceptionally low proliferation rate (0.015%) and CD11c+ classical DCs (0.08%) (**Fig. 3C**).

**Fig. 3.**
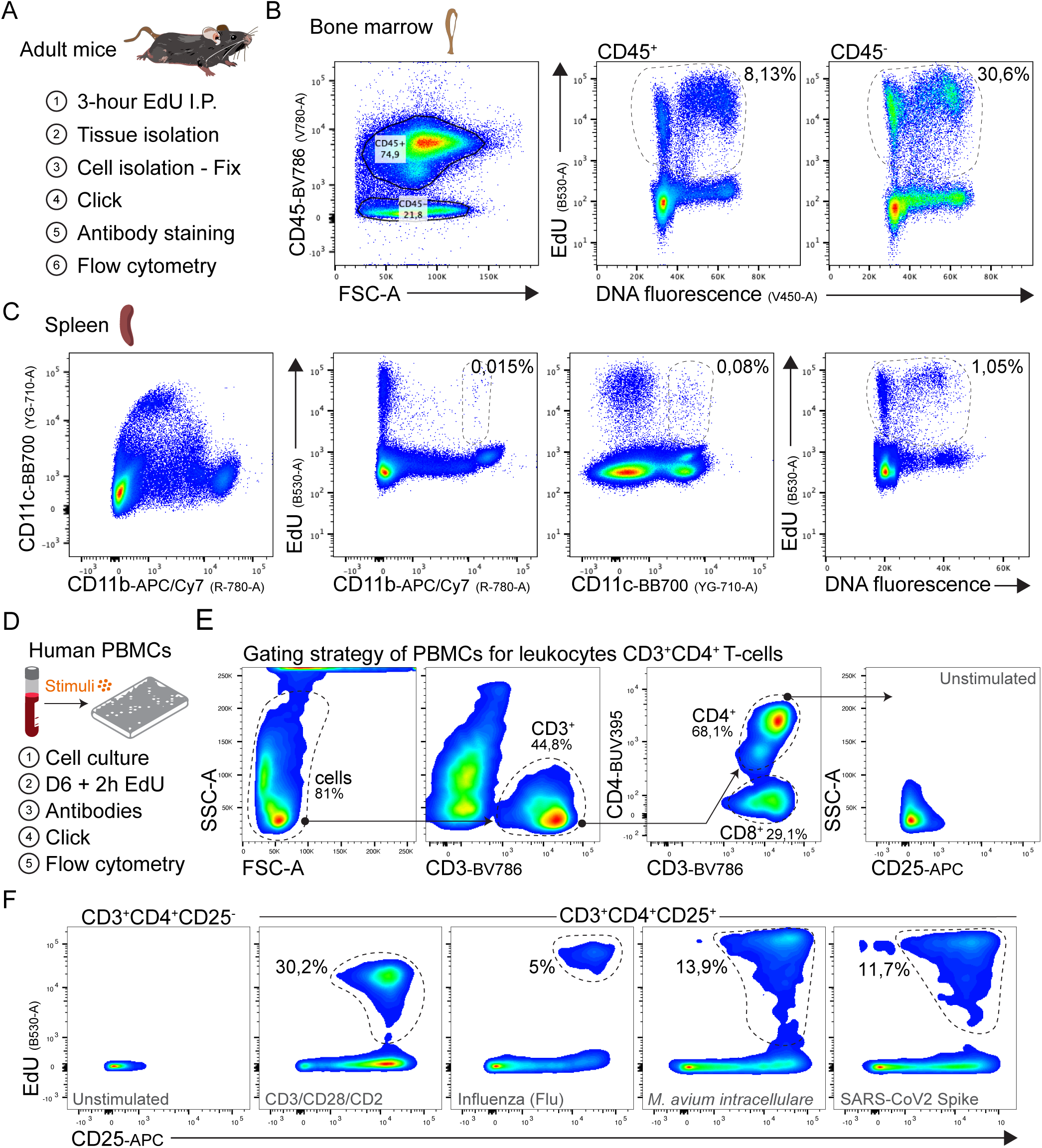
Compatibility of OpenEMMU for multi-parametric analysis of DNA replication in mouse and human immune cells. (A) Outline of adult mice EdU injection and OpenEMMU protocol for studying DNA synthesis in various tissues and cells. (B) Flow cytometry of EdU signal using isolated bone marrow cells (*n* = 3). (C) Flow cytometry of EdU signal using isolated splenocytes, highlighting percentages of rare DNA replicating cells (CD11b+ and CD11c+) (*n* = 3). (D) Outline of human PBMCs evaluation for EdU incorporation and responses to various stimuli. (E) Gating strategy employed to detect DNA replication in activated human T-cells. The dashed lines indicate the selected gates. (F) Cell stimulation with activating antibodies or exposure to infectious disease-relevant immune stimulants showing the quantification of EdU+ DNA replicating CD3^+^CD4^+^CD25^+^ cells (dashed lines) (*n* = 2).

To demonstrate OpenEMMU’s clinical relevance while extending its multiplexing scope, we next evaluated human T-cell activation and CD3+CD4+ T-cell proliferation using peripheral blood mononuclear cells (PBMC) cultures. CellTrace^TM^ Violet (CTV) labeled PBMC were cultured for 6 days using separate negative control cultures (medium only), positive control cultures with optimal T-cell polyclonal antibody activation (CD3/CD28/CD2), as well as other cultures for proliferative responses to various recall antigens from infectious agents, including (i) influenza vaccine antigen; *M. avium intracellulare* lysate; and (iii) SARS-CoV-2 spike protein, respectively (**Fig. 3D-F**). In all tested conditions, proliferating EdU-labeled CD3+CD4+CD25+CTVdim T cells exhibited DNA replication in response to canonical CD3/CD28/CD2 stimulation and recall antigens, albeit at varying rates (**Fig. 3F**). Our findings support earlier observations, confirming OpenEMMU’s multiplexing capabilities with various fluorochrome-conjugated antibodies. The simultaneous detection of EdU fluorescence alongside CD45, CD11b, CD11c, T-cell lineage markers (CD3, CD4, CD8), T-cell activation marker CD25, and cell division marker CTV, suggests that EdU-labeled, OpenEMMU-processed cells can be used to investigate additional cell markers across various cell types—a capability previously limited with commercial kits.

### 3D imaging of DNA replication and cycling cells in adult mice organs and IBEX multiplexing

Following the flow cytometric evaluation of OpenEMMU’s performance using dissociated single cells isolated from mice injected with EdU for 4 hours, as described above, we applied the method to label cells in 3D mouse tissues, focusing first on the stem cell niche within the adult small intestine as a homeostatic proliferative and regenerative model. To achieve this, we combined OpenEMMU with ethyl cinnamate (ECi) tissue clearing. Laser confocal imaging of ECi-cleared ileum showed extensive EdU labeling of DNA-replicating cells across a region ∼100 µm in depth (**Fig. 4A, Fig. S6B**). This enabled targeted confocal microscopy at different focal planes to visualize specialized regions within the adult stem cell niche, such as the stem cell crypt, villus, WGA-enriched Paneth cells, and highly DNA replicating transient-amplifying cells (**Fig. 4A**). We confirmed that either Saponin or Triton X-100-containing cell permeabilization/wash buffers could be used successfully in the OpenEMMU assay (**Fig. S6C**). Lateral or top-view 3D imaging can be used for visualizing DNA replication and cycling cells (**Fig. 4A, Fig. S6B-D**). Four hours of intraperitoneal EdU administration allowed for the successful identification of not only DNA-replicating cells but also cells that had progressed through the cell cycle (S and G2 phase) and reached mitosis, as judged by condensed chromosomes in metaphase (**Fig. 4A**).

**Fig. 4.**
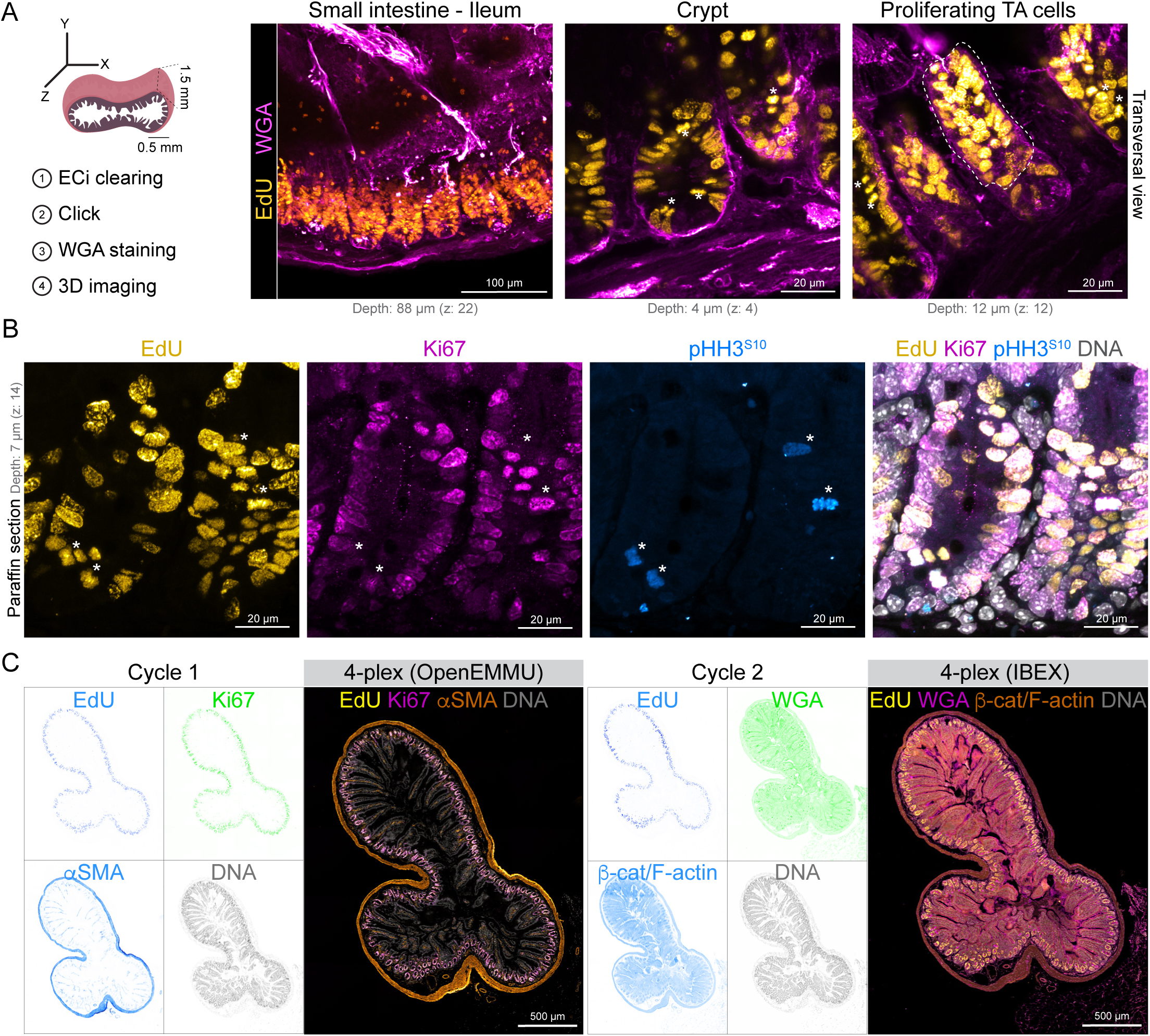
OpenEMMU for multiplexed imaging and DNA replication studies in the adult intestinal stem cell niche. (A) OpenEMMU compatibility with ethyl cinnamate tissue clearing and 3D confocal laser imaging, during analysis of DNA synthesis in the small intestine (ileum) of adult mice injected with EdU for 4 hours. Several proliferating cell regions are observed, including the adult stem cell crypt and the transient amplifying (TA) cells in the villi, highlighted by the dotted line. (B) FFPE-processed ileum section and confocal laser imaging of EdU in DNA-replicating cells, along with Ki67 and p-HH3^S10^ that are EdU positive, visualized by *z*-stack confocal microscopy. (C) FFPE-processed ileum section, visualized using tiling confocal microscopy. The section underwent two cycles of 4-plex staining, with OpenEMMU used in the first cycle and IBEX applied in the second (*n* = 3).

To further validate cell cycle progression assessment and immunolabelling compatibility of OpenEMMU, we stained formalin-fixed paraffin-embedded (FFPE) ileum sections using an optimized protocol for FFPE, deparaffinization, and antigen retrieval (Zaqout, Becker, and Kaindl, 2020) (**Fig. 4B, Fig. S6A**). *In vivo* EdU labeling was extensively observed along the gut stem cell niche in highly DNA-replicating cells with successful co-immunostaining of a proportion of cells with Ki67 (a nucleolar protein and proliferative capacity marker) and p-HH3^S10^ (a mitotic marker) (**Fig. 4B**). DNA labeling was also achieved. We confirmed that only a few EdU-labeled cells had reached mitosis (p-HH3^S10+^) during the labeling period and observed that, in several cases, Ki67 staining was weaker than EdU staining for actively cycling cells (**Fig. 4B**).

To enable multiplexed imaging, we employed OpenEMMU followed by iterative bleaching extends multiplexity (IBEX) (Radtke et al., 2020). IBEX is an open-source iterative immunolabelling and chemical bleaching technique that offers a low-cost and accessible approach using borohydride derivatives, requiring only basic laboratory skills (Radtke et al., 2022). To assess the compatibility of OpenEMMU with this relatively novel multiplexing method, we first labeled FFPE ileum sections for Ki67 (R667), αSMA (Cy3), EdU (AZDye 488 Azide), and DNA (Vybrant Violet) during the first imaging cycle (**Fig. 4C**). Following the completion of this round (cycle 1), we applied LiBH□ (1 mg/mL) to inactivate the fluorophores and proceeded with a second staining cycle (cycle 2), where we stained for WGA (CF640R) and β-catenin (PE) together with F-actin (Flash Phalloidin 594) (**Fig. 4C**). Our observations revealed that EdU (AZDye 488 Azide) and Vybrant Violet (DNA staining) exhibited significant resistance to LiBH□ bleaching, as strong signals persisted even after two rounds of LiBH□ incubation. Consequently, re-labeling for EdU or DNA was unnecessary. A third immunolabelling cycle (cycle 3) was subsequently performed, staining for E-cadherin (A555) and COL1A1 (A555) simultaneously, and BODIPY 630/650 (**Fig. S6E**). We noted that CF640R (WGA) also demonstrated partial resistance to LiBH-mediated fluorophore inactivation, requiring longer bleaching. These findings show that OpenEMMU can be multiplexed to capture the *in-situ* biology of complex tissues with precision and spatial resolution.

### Mapping DNA replication and cell proliferation in developing mouse embryos and organs

To further investigate the applicability of OpenEMMU to studying DNA replication and cell proliferation *in vivo*, we assessed (i) its ability to identify proliferative foci across other developing organs and tissues; (ii) the preservation of fine and fragile embryonic tissue structures in FFPE mouse embryos after deparaffinization, OpenEMMU click chemistry and antibody staining; and its compatibility with complex 3D immunostaining using fluorochrome-conjugated and non-conjugated antibodies.

To label developing embryos, pregnant females were administered a 3-hour pulse of EdU. Using anti-Ki67, αSMA-Cy3, and p-HH3^S10^ immunohistochemistry, we imaged whole mouse embryo sections at E14.5 comprehensively, capturing the characteristic organ localization and organization using tile confocal microscopy and the Thunder Imager epifluorescence system for accelerated image acquisition (**Fig. 5, Fig. S7**). We observed EdU-positive cells with distinct nuclear labeling abundance, with different tissues and organs exhibiting varying degrees of DNA-replicating cells. Whereas some cells in specific organs and tissues such as the liver, lungs, and lateral ventricle (LV) of the brain were highly proliferative (higher EdU uptake), others, like those in the midbrain region, the developing ureteric bud region of the kidneys, and to a lesser extent developing skeletal muscles and heart, had a much-reduced proliferative rate (lower EdU uptake) (**Fig. 5A and B, Fig. S7A**). Next, we labeled earlier developing embryos at E11.5 with EdU for 3 hours. OpenEMMU enabled precise visualization of EdU+ cells alongside co-staining for Ki67, αSMA-Cy3, p-HH3^S10^, and DNA across various developing organs and tissues (**Fig. S7B and C**). As evident in highly magnified regions, finer subcellular structures were well preserved throughout the deparaffinization and OpenEMMU procedures.

**Fig. 5.**
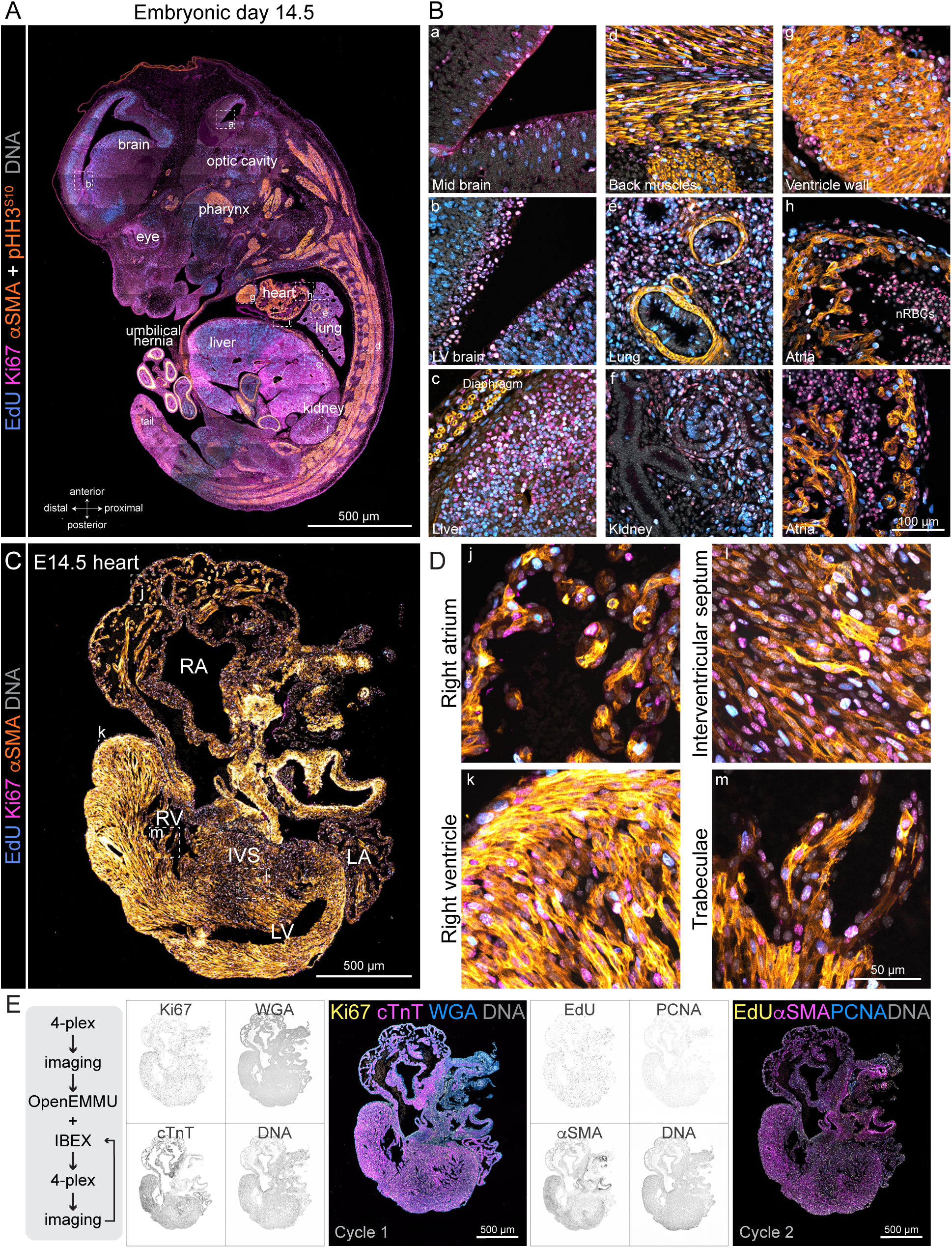
Embryo and organ-specific analysis of DNA replication and mitosis in developing mouse embryos. (A) FFPE-processed embryonic day E14.5 mouse embryo and confocal tile imaging of EdU signal in DNA replicating cells multiplexed with Ki67, αSMA, phospho-histone H3^S10^, and DNA labeling. αSMA immunolabelling depicts the developing musculature, heart cardiomyocytes, as well as smooth muscle cells in multiple organs and tissues (*n* = 3). Insets (B) show the magnification of the boxed areas using *z*-stack confocal microscopy (a-i). (C) FFPE-processed E14.5 developing heart and confocal tile imaging of EdU signal in DNA replicating cells multiplexed with Ki67, αSMA, and DNA labeling (*n* = 2). αSMA immunolabelling depicts the developing musculature, heart cardiomyocytes, and smooth muscle cells in multiple cardiac regions. Insets (D) show the magnification of the boxed areas using *z*-stack confocal microscopy (j-m). (E) E14.5 heart section underwent two 4-plex staining rounds: first with conjugated antibodies and WGA, then with OpenEMMU and IBEX (*n* = 3).

The adult four-chambered mammalian heart is the first organ to form and function in the embryo. To further elucidate DNA replication and cell proliferation in the developing mouse heart, we applied OpenEMMU. By E14.5, most cardiac structures have successfully developed, including the upper chambers (atria), distinct left and right ventricles, interventricular septum, transient outflow tract, and elements of the cardiac conduction system. Specific heart tissues and regions can be distinguished, such as the myocardium, epicardium, endocardium, coronary vasculature, trabeculae, and cushion tissues (**Fig. 5C and D**). Using OpenEMMU in combination with immunohistochemistry against Ki67 and αSMA, we were able to focus and distinguish DNA-replicating and cycling cells in the developing heart, enabling the faithful detection of DNA synthesis in cardiomyocytes and non-cardiomyocyte stromal cells (**Fig. 5C and D**).

We then assessed whether OpenEMMU could be applied following an initial cycle of 3-plex immunostaining with DNA staining, followed by imaging, IBEX, OpenEMMU, and two additional IBEX cycles (**Fig. 5E**). We successfully stained for Ki67-Vio®-B515, WGA-CF®555, cTnT-AF®647, EdU-AZDye 488, PCNA-AF®647, αSMA-Cy3, COL1A1-AF®647, MLC2v-PE, MLC2a-APC, and p-HH3^S10^-PE, allowing the visualization of multiple spatial features within the developing heart (**Fig. 5E**). Notably, Ki67 labeling was extensive, similar to PCNA, while EdU coverage was nuclear and more limited. Only a few rare mitotic cells (p-HH3^S10^) were observed, and these were primarily within the cardiac ventricles (MLC2v+) (**Fig. 5E and Fig. S7D**). MLC2a specifically marked the developing left and right atria, whereas WGA staining was found throughout, strongest within the epicardium (**Fig. 5E and Fig. S7D**). Thus, OpenEMMU can be effectively applied across ≥4 staining cycles, both before (**Fig. 4C and Fig. S7D**) and after IBEX (**Fig. 4C and Fig. 5E**), maintaining tissue and signal integrity.

To investigate whether OpenEMMU could be used to profile individual cells and their cell cycle accurately and quantitatively during heart development, we first isolated E14-developing hearts and performed immunolabeling against PDGFRα and CD31, along with detecting EdU+ cells by flow cytometry (**Fig. 6A**). Visualization of these distinct populations of cells showed that approximately 13% of fibroblasts and 15% of endothelial cells were stained with a 3-hour EdU pulse (**Fig. 6B**). Confocal imaging-based assessment of total dissociated WGA-stained cells demonstrated that approximately 12% were EdU+ (**Fig. S8A-C)**. We were able to profile both stromal cells and cardiomyocytes expressing cardiac muscle Troponin T (TNNT2, also known as cTnT), which exhibited strong EdU labeling at this developmental stage (**Fig. S8D**). Together, these results establish that OpenEMMU is compatible with a spectrum of staining procedures for which existing methods are not fully optimized.

**Fig. 6.**
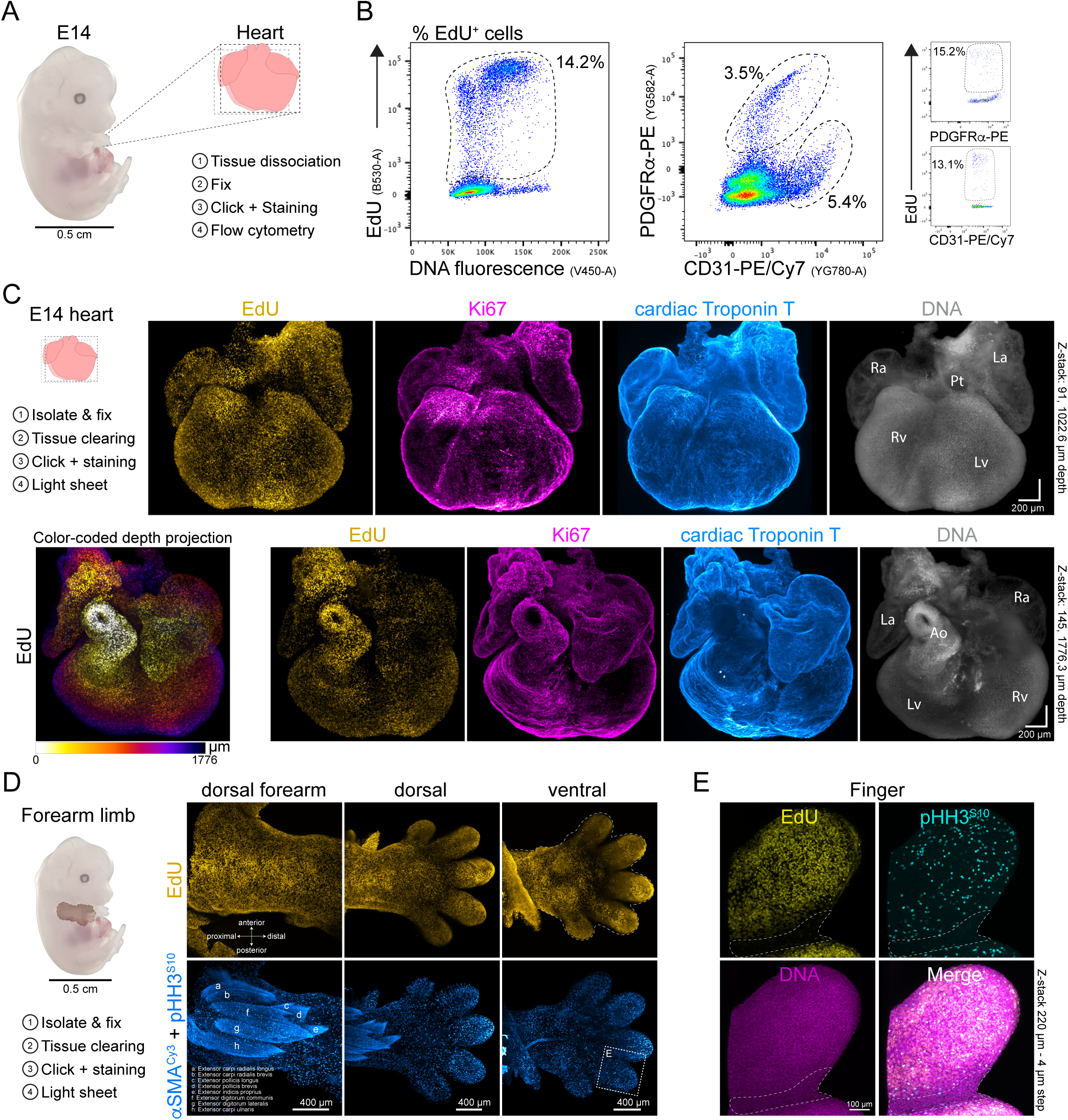
Application of OpenEMMU to complex developing organs and tissues. (A) Diagram of an embryonic day 14 (E14) mouse embryo showing the steps involved in isolating the heart and the OpenEMMU protocol. (B) Flow cytometry results of isolated total heart cells with EdU staining along DNA content (left graph), demonstrating its multiplexing capability with CD31-PE/Cy7 and PDGFRα-PE. The relative proportion of EdU+ cells in both cell populations is depicted (*n* = 3). (C) Light-sheet fluorescence imaging of EdU multiplexed with Ki67 and cardiac Troponin T, along with DNA labeling in an E14 heart, shown from a front view (top panels) (*n* = 4). A top view of the same heart is provided, with EdU labeling color-coded by depth projection. Ra, right atrium; La, left atrium; Pt, pulmonary track; Rv, right ventricle; Lv, left ventricle; Ao, aorta. (D) Light-sheet imaging of EdU multiplexed with αSMACy3 and p-HH3^S10^-A594 immunolabeling, along with DNA staining of an E14 forelimb, shown from dorsal and ventral views (*n* = 3). (E) Inset of (D, ventral side of the digit) showing a magnified view of the boxed area using Z-stack confocal microscopy. The dotted lines indicate the interdigital necrotic zone, where cellular DNA synthesis and mitosis decrease dramatically.

### Mapping DNA replication and cell proliferation in whole-organ 3D fluorescence imaging

Due to the limitations of commercial EdU kits in permeabilizing and penetrating complex 3D organs and structures, sensitive 3D imaging is often limited. We sought to investigate the applicability of OpenEMMU for detecting DNA-replicating cells in whole developing organs, including the heart and limbs. Using DEEP-Clear tissue clearing (Pende et al., 2020) and light-sheet fluorescence microscopy (LSFM) on a developing mouse heart at E14, we achieved 3D visualization of major mammalian cardiac structures through immunohistochemistry against cardiac Troponin T, highlighting extensive EdU labeling at this developmental stage (**Fig. 6C**). Notably, EdU labeling outperformed Ki67 in detecting cells in the S-phase of the cell cycle, as areas containing EdU-labeled nuclei were often only partially positive for Ki67 **(Fig. 6C**). These differences can be attributed to (i) the significantly enhanced brightness of EdU detection using OpenEMMU, and (ii) the highly heterogeneous dynamics of Ki67 protein abundance throughout the cell cycle (e.g., G0/G1 cells) (Miller et al., 2018).

To further explore OpenEMMU 3D imaging capabilities, we evaluated EdU labeling in an E14 (TS21 to TS22+) forearm limb using LSFM (**Fig. 6D**). We observed extensive DNA synthesis, as indicated by EdU staining, and co-detected rounded, chromatin-dense mitotic cells using a PE-conjugated antibody against p-HH3^S10^ (**Fig. 6D, Fig. S8E**). OpenEMMU, in combination with αSMA-Cy3 immunostaining, enabled the visualization of distinct developing extensor muscle groups of the forearm with myofiber resolution (**Fig. 6D, Fig. S8E**) (Besse et al., 2020). EdU-labeled cells and mitotic cells could also be distinguished with single-cell resolution using not only LSFM but also confocal microscopy (**Fig. 6D**, **Fig. 6E**). Notably, a detailed YZ-axis view of light-sheet images of the developing forearm extensor muscles showed labeling at imaging depths of over 150 µm (**Fig. S8F**). These results demonstrate the extended applicability and scalability of OpenEMMU to more complex organs and tissues, preserving morphological features and staining quality.

### Profiling DNA replication and cell proliferation in 3D self-organizing cardiac organoids

iPSC and embryonic stem cell-derived organoids have revolutionized our understanding of tissue and organ morphogenesis, as well as cell fate and behavior (Cerneckis et al., 2024). Organoids serve as invaluable research platforms for investigating cell specification, organ growth, and maturation within complex and physiologically relevant 3D environments, heralding significant advancements in biomedical research. Given the current challenges in accurately modeling early human heart development, 3D cardiac organoids derived from human iPSCs present a promising platform for studying cardiac development and congenital heart disease (Cho et al., 2022; Zhang and Wu, 2024), which affects approximately 1% of newborns worldwide (Jenkins et al., 2007).

We evaluated OpenEMMU’s performance in labeling and visualizing self-organized hiPSC-derived cardiac organoids (hCOs) at days 12 and 19 of differentiation and early maturation (**Fig. 7**), observing effective staining of proliferative cells within beating organoids, albeit at varying intensities across distinct regions (**Fig. 7B, Fig. S9**). Overall, we observed that the majority of internal regions exhibited a weak EdU label. This may be attributed to slower cell cycle progression in comparison to cells nearer the hCO’s periphery (media boundary), likely due to cell differentiation, tissue maturation, or diminished tissue perfusion (**Fig. 7B, Fig. S9**). When co-staining for the NKX2-5 cardiac transcription factor, we noted that several NKX2-5-expressing cardiomyocytes were also positive for EdU, while others were not (**Fig. 7B**). These cells were also embedded within cardiac Troponin T-expressing regions, confirming their cardiac and sarcomeric contractile identity (**Fig. 7B, Fig. S9**).

**Fig. 7.**
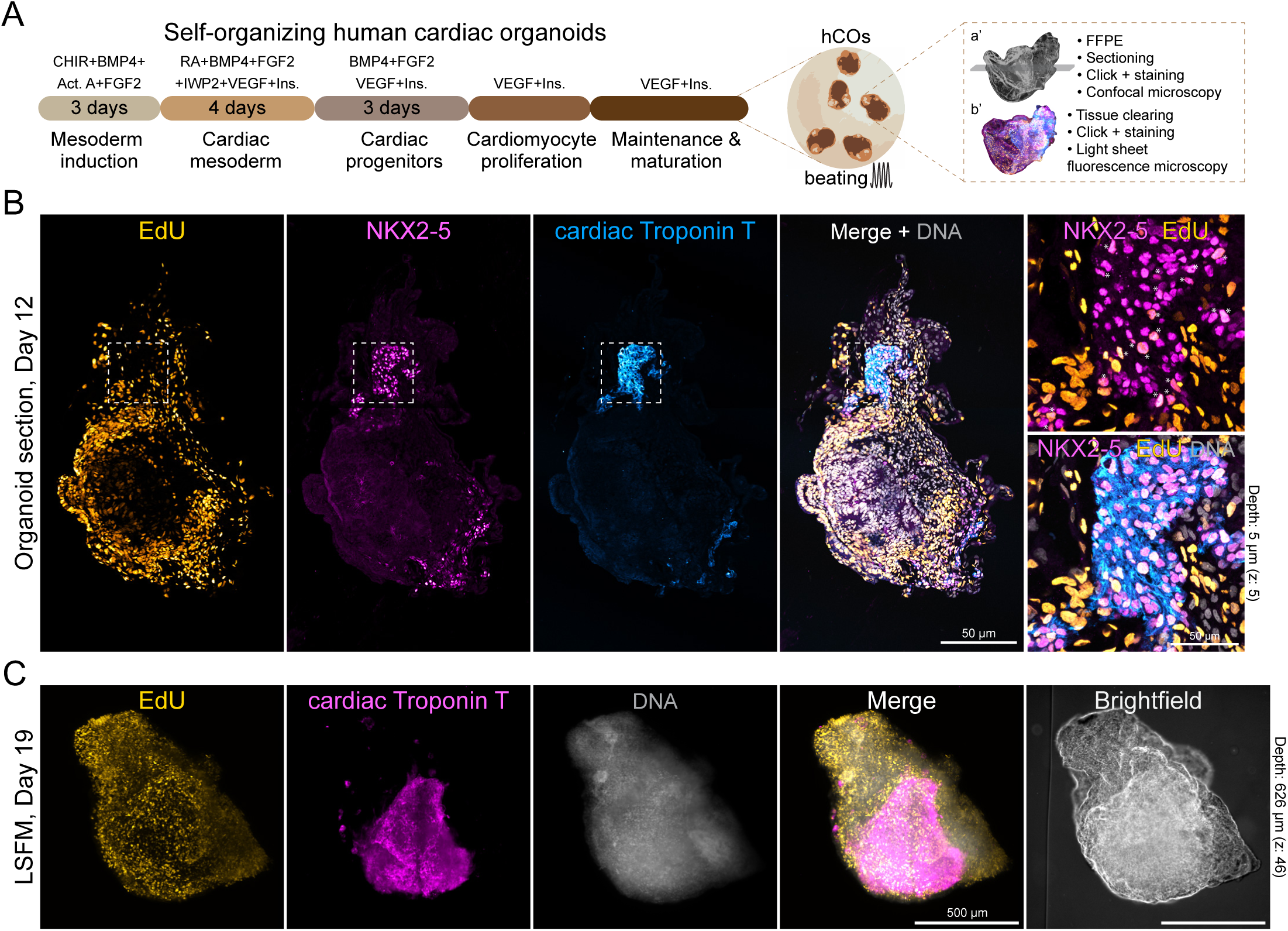
DNA replication in self-organizing 3D human cardiac organoids. (A) Protocol outline to generate self-organizing and beating hiPSC-derived human cardiac organoids (hCO), showing two distinct imaging strategies: FFPE organoid confocal imaging (a’) and 3D light sheet microscopy (b’). (B) FFPE-processed day 12 organoid and confocal tile imaging of EdU fluorescence in DNA replicating cells (*n* = 6). NKX2-5 and cTnT expressing cardiac progenitors are highlighted in the boxed area. NKX2-5 and EdU double-positive cells are also shown (*, magnified box). (C) Whole-mount light sheet microscopy (LSFM) 3D visualization of a hCO at day 19 showing EdU positive cells and cTnT expressing cells. DNA-labelled nuclei are also shown (*n* = 6).

Because of the challenges associated with using FFPE for relatively miniature and fragile organoids, we optimized an OpenEMMU labeling and organoid-clearing strategy to visualize complex 3D structures within hCOs better using LSFM. As previously observed, OpenEMMU was compatible with the staining of 3D structures. Resulting images helped identify cardiac Troponin T+ regions with depths exceeding 500 µm (**Fig. 7C**). Additionally, strong EdU staining was also observed in the non-cardiac areas (**Fig. 7C**). Together, our results demonstrate that OpenEMMU, in combination with immunofluorescence, can be used to gain a deeper understanding of DNA replication and cell proliferation in human developing organs and tissues. Moreover, it is effective in organoid-derived platforms, with the potential to offer insights into cell proliferation in both health and disease.

### Analysis of DNA replication in Zebrafish larvae

Zebrafish are known for their small size, rapid external development, imaging clarity, and low-cost husbandry, making them a valuable model for vertebrate development and human research (Veldman, M. B., & Lin, 2008; Teame et al., 2019). Zebrafish embryos and larvae have been effectively utilized to investigate the biological impact of a variety of compounds, drugs, and chemicals, on different organs and tissues (Bauer, Mally, Liedtke, 2021; Cassar et al., 2020). They complement mammalian models, offering advantages for large-scale genetic studies and rapid gene function analyses.

Following a 2-hour EdU immersion protocol of *Danio rerio* larvae 5 days post-fertilization (dpf), OpenEMMU facilitated a detailed anteroposterior overview of DNA synthesis using FFPE samples, in combination with αSMA immunostaining and DNA staining (**Fig. 8A and B, Fig. S10A-C**). By 5 dpf, zebrafish larvae are free-swimming and independent feeders, although many of their organs and structures, such as the caudal and pectoral fins, developing musculature, brain, eyes, and jaws, continue to develop (Kimmel et al., 1995). EdU-labeled cells were detectable in highly proliferative regions across the developing larvae, including the ciliary marginal zone (CMZ) in the developing teleost retina (Fischer et al., 2013), the developing myotome, and notochord (**Fig. 8B**). Sporadic mitotic chromosomes were identified using OpenEMMU (**Fig. 8B**, magnified regions), demonstrating that EdU-labelled cells can progress from the S phase to the G2 and M phases of the cell cycle in 5dfp developing larvae. These results were obtained using either confocal laser microscopy or the Thunder Imager epifluorescence system for faster data acquisition (**Fig. 8B, Fig. S10C**).

**Fig. 8.**
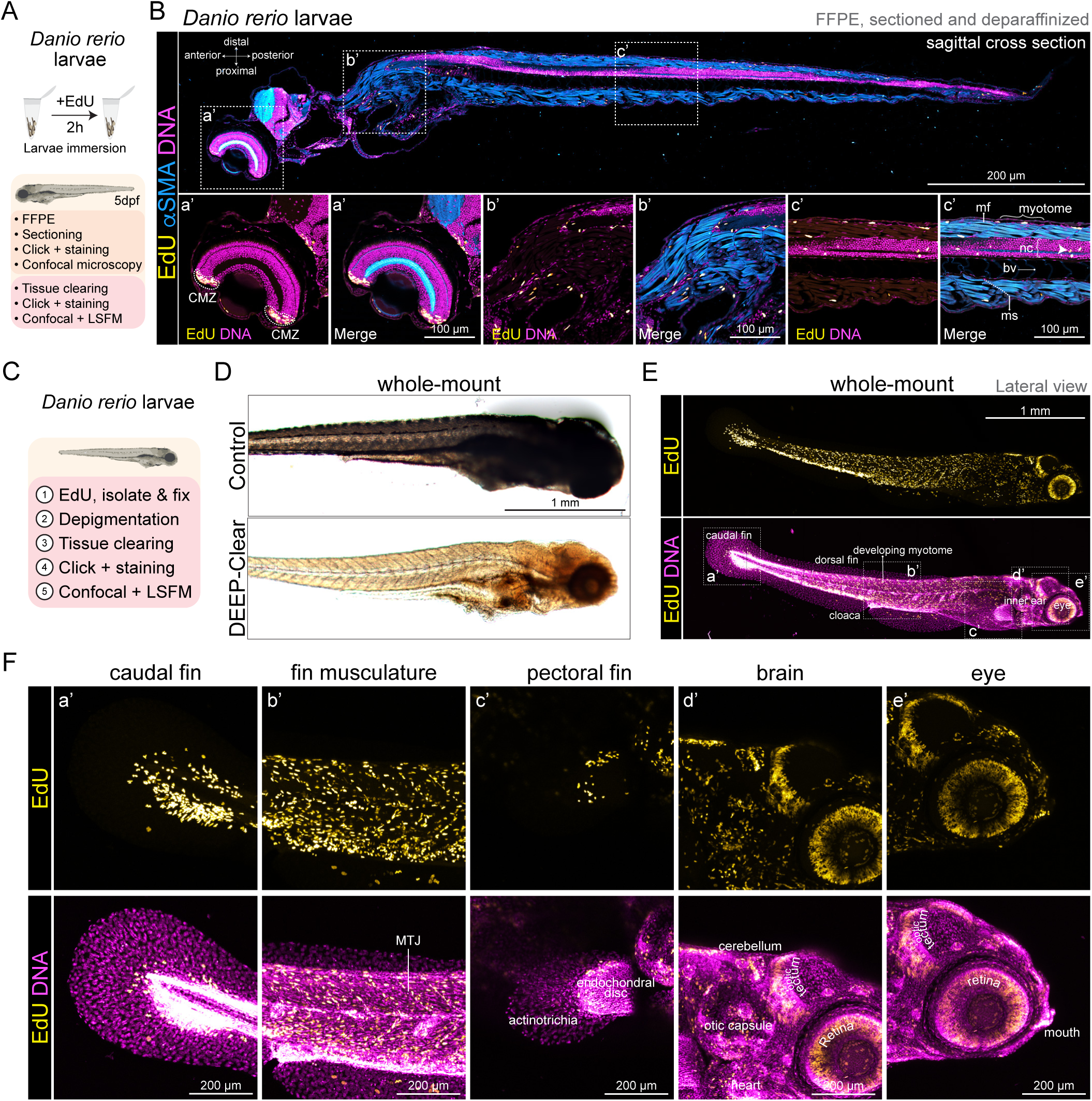
Analysis of DNA replication in whole zebrafish larvae. (A) *Danio rerio* 5dpf larvae immersion protocol aimed at labeling DNA synthesis with EdU for 2h using FFPE (B) or whole-mount staining (D-F). (B) FFPE-processed larva sagittal cross section and confocal tile imaging of EdU signal in DNA replicating cells (*n* = 3). αSMA-Cy3 immunolabeling depicts the developing musculature and growing myotomes. Insets (below) show the magnification of the boxed areas (a’-c’). Arrowhead shows a mitotic event. (C and D) Depigmentation and DEEP-Clear protocol allow whole-mount visualization of growing larvae (*n* = 6). (E and F) EdU and DNA labeling revealing highly proliferative cells, visualized by tiling (E) and Z-stack (F) confocal microscopy. Insets (F) show the magnification of the boxed areas (a’-e’). CMZ, ciliary marginal zone; MTJ, myotendinous junction; nc, notochord; ms, myoseptum; mf, myofiber; bv, blood vessels.

Having confirmed OpenEMMU’s efficacy in 3D staining of whole organs in mice and iPSC-derived organoids, we combined *in vivo* labeling of nascent DNA using whole-organism tissue clearing (i.e., DEEP-Clear (Pende et al., 2020)) to comprehensively visualize DNA replication activity in 5dpf zebrafish larvae (**Fig. 8C**). Depigmentation and 3D tissue clearing enabled whole-mount visualization of the 5dpf zebrafish larvae and facilitated OpenEMMU for effective 3D nascent DNA visualization across the entire organism (**Fig. 8D and E, Fig. S10D**) and at different focal planes, as observed with either confocal laser microscopy or LSFM (**Fig. 8E, Fig. S10E and F**). EdU-labeled cells were extensively observed in the growing caudal fin and the distal portion of the endochondral disc of the pectoral fin (**Fig. 8F**). They were also present across the developing myotome, including the myotendinous junctions (**Fig. 8F**). Additionally, these cells were found in a cellular structure on the lateral surface of the myotome known as the external cell layer (ECL) (**Fig. 8F**), which is responsible for generating stem and progenitor cells for larval muscle growth (Nguyen et al., 2017). Distinct regions of the larvae’s head, such as the cerebellum, the optic tectum, and the otic capsule, also showed substantial EdU labeling (**Fig. 8F**). Additionally, extensive DNA replication was observed in the developing retina and mouth **(Fig. 8F, Fig. S10F**). Finally, we visualized regenerative adult zebrafish ventricular resections using OpenEMMU. EdU-labeled cells were observed throughout the trabeculated myocardium, with enrichment in the regenerating ventricular apex, as anticipated (**Fig. S10G**). We demonstrate here that OpenEMMU further enhances the utility of zebrafish for studying DNA replication and cell proliferation in both physiological and pathophysiological conditions. OpenEMMU is therefore potentially well-suited for studying tissue and organ regeneration through EdU uptake experiments in biomedically relevant animal models, including zebrafish.

## DISCUSSION

### Development and optimization of clickable OpenEMMU to measure DNA replication and cell-cycle progression

We report the development and optimization of a robust and open-source toolkit for DNA replication profiling and cell cycle measurement using CuAAC click chemistry. OpenEMMU facilitates efficient detection of EdU incorporation in cultured cells, tissues, organs, and entire organisms using defined and cost-effective click chemistry reagents, significantly advancing the detection and analysis of DNA synthesis across various cell types and models, including human cells. We sequentially analyzed the influence of each chemical reagent to determine the conditions required for optimal labeling efficiency, as well as the quality and intensity of EdU fluorescence. Our systematic approach in refining the EdU click reaction demonstrates that OpenEMMU surpasses the performance of commercially available kits, particularly in terms of cost-effectiveness, versatility, and signal detection.

The key optimizations, including the determination of 800 µM as the crucial reagent concentration for CuSO□ in the OpenEMMU assay, underscore the importance of fine-tuning reaction components to achieve high-efficiency labeling and superior signal-to-noise ratio. This adjustment, along with the in-house optimization of the permeabilization/wash buffer, serum, L-ascorbic acid, and picolyl azide concentrations, has resulted in a highly sensitive and specific custom assay capable of accurately distinguishing between various phases of the cell cycle and measuring DNA replication in different cellular contexts. The substantial improvement in EdU signal intensity (∼10-fold) compared to the Invitrogen commercial kit further validates the efficacy of OpenEMMU, particularly in scenarios requiring the co-detection of DNA synthesis and other cellular markers, PE-conjugated antibodies, IBEX-mediated imaging multiplexing, and complex 3D organs and organisms.

### The broad applicability of OpenEMMU across different cell types, organs, and tissues

OpenEMMU’s performance and its ability to sensitively detect DNA replication across a spectrum of cellular and molecular contexts, from proliferative hiPSCs, C2C12, C3H10T1/2, pancreatic and breast cancer cell lines, and growth factor-stimulated cell proliferation to low-replicating hiPSC-derived cardiomyocytes, activated human T cells, and *in vivo* murine cells (e.g., small intestine, bone marrow, and spleen), showcases its broad applicability, robustness and adaptability. The method’s compatibility with fluorescence imaging, flow cytometry, and FACS should enhance researchers’ ability to dissect cell cycle dynamics in complex biological problems with high precision and combine OpenEMMU with genomics and spatial-’omics approaches. Moreover, the ability to multiplex EdU detection with various immunolabeling techniques, including the detection of cell surface markers and intracellular and nuclear proteins, enhances its utility in studies of cell identity, cell differentiation, organismal development, and disease.

One of OpenEMMU’s most compelling advantages is its resource efficiency. By utilizing commonly available laboratory reagents, OpenEMMU offers a highly economical alternative to commercial kits, costing approximately USD 0.05 per 1 mL of clickable solution—142 times cheaper than Invitrogen and 77 times cheaper than VectorLabs kits. This makes sophisticated DNA replication studies more accessible to diverse scientific communities. We have provided step-by-step imaging and flow cytometry protocols to guide the OpenEMMU user through the various protocols, making it labor-efficient. Its affordability and user-friendliness do not come at the expense of performance; rather, OpenEMMU outperforms commercial kits in several key aspects, including signal intensity, compatibility with PE-conjugated and non-conjugated antibodies, and the ability to label a wide range of cell types and complex 3D tissues and structures. Our open-source philosophy should allow for further advancements. Its compatibility with novel iterative labeling methods, such as IBEX, further enhances its utility for comprehensive biological studies, positioning it to revolutionize multi-target fluorescence research.

### OpenEMMU in multiplexed immunological and human T-cell activation and proliferation

Our findings further underscore OpenEMMU’s utility as an adaptable tool for multiplexed analysis in immunological studies, such as in the context of T-cell activation and proliferation. Notably, this is the first known application of the EdU thymidine analog to assess human T-cell activation and DNA replication in response to several physiologically relevant stimuli (Alvarez et al., 2020; Ganesan et al., 2023), including common anti-CD3/anti-CD28/anti-CD2 polyclonal activators, influenza vaccine, *M. avium intracellulare*, and recombinant trimeric SARS-CoV-2 spike protein. OpenEMMU’s ability to simultaneously detect DNA replication and a variety of surface markers in human T cells creates new opportunities for in-depth investigations into lymphocyte immune responses under typical and pathogenic challenges. Looking forward, OpenEMMU should help unveil new perspectives on immune cell populations and functional states in both health and disease, such as within tumor microenvironments or autoimmune conditions.

### OpenEMMU for whole-organ, organoid, and whole-organism 3D fluorescence imaging of nascent DNA

The application of OpenEMMU in 3D tissue imaging and whole-organ studies—such as mouse embryonic development, heart and forelimb development, and zebrafish—opens new avenues for exploring DNA replication in complex biological systems. This is achieved using cutting-edge technologies, including tissue clearing (Susaki & Ueda, 2016; Richardson et al., 2021, **Materials and Methods**) and advanced imaging techniques like light-sheet fluorescence microscopy (Stelzer et al., 2021). The ability to map DNA replication in 3D in developing organs, particularly in spatially challenging contexts like heart development, whole-mount zebrafish larvae, and self-organizing cardiac organoids, is a significant advancement, and this capability will facilitate understanding of organ and tissue regeneration. OpenEMMU allowed us to distinguish varying degrees of DNA replication, revealing significant differences and proliferative foci across multiple organs and cell types. The 3D mapping of DNA replication activity as it correlates with cell division rates, transcriptional activity, and metabolic processes, will deepen our understanding of the complexities of growth, function, and morphology in developing organisms.

OpenEMMU has proven to be fast, efficient, affordable, and scalable. These advances are expected to significantly influence future research on cell-cycle progression in various cell types through cost-effective, open-source assays, including the validation and extension of previous findings using thymidine analog uptake (e.g., BrdU, EdU). CuAAC click chemistry also has multiple biomedical and non-biomedical applications, including radiochemistry (Bauer et al., 2023), nucleic acid (Fantoni et al., 2021), RNA (Tamura et al., 2024), glycan (Zhang & Zhang, 2013), lipid (Jamecna & Höglinger, 2024), in vivo drug binding (Pang et al., 2022), and protein biology (Parker & Pratt, 2020). Thus, open-source and reliable click chemistry methods are benefiting the development of custom assays and novel techniques in biotechnology, biomedicine, biochemistry, nanotechnology, and materials science.

In summary, our study introduces OpenEMMU, an open-source toolkit for DNA replication profiling and cell cycle analysis using CuAAC click chemistry. This dual modality enhances the sensitivity and resolution of DNA synthesis assessments and simplifies experimental workflows, representing a substantial advancement over existing protocols. By optimizing key reagents such as CuSO□, ascorbic acid, and picolyl azide, we achieved improved and more reliable CuAAC click chemistry compared to standard commercial kits. We successfully integrated OpenEMMU with tissue depigmentation, custom cell permeabilization, blocking protocols, multiplex imaging using IBEX, and advanced tissue-clearing techniques like DEEP-Clear. Validated in various 2D cell cultures and complex 3D structures, OpenEMMU offers reproducible workflows for efficient DNA replication characterization without the need for commercial kits. This platform enhances scalability, significantly reduces costs, and supports researchers across diverse models and systems. OpenEMMU has broad applications in cancer biology, immunology, stem cell and developmental biology, pathophysiology, and disease modeling, with the potential to become a standard technique for DNA replication and cell cycle research.

## LIMITATIONS OF THE STUDY

Our research concentrated on utilizing Cu(I)-Catalyzed Azide−Alkyne Cycloaddition (CuAAC) click chemistry to monitor and assess DNA replication. It is crucial to acknowledge that other click chemistry reactions may demonstrate different efficiencies and labeling characteristics. Additionally, biases or technical limitations could arise from variations in experimental design, such as pulse-chase experiments, labeling duration, and the reagents employed. Therefore, significant new insights into cell cycle dynamics will necessitate validation across multiple laboratories and through diverse methodologies.

## MATERIALS AND METHODS

### Ethics statement and Mouse strain

Wild type [WT, Inbred C57BL/6J] (Jackson Laboratory; 000664) mice were bred and housed under pathogen-free conditions in the BioCORE facility (Victor Chang Cardiac Research Institute, Sydney, Australia). All experimental procedures were approved by the Garvan Institute/St. Vincent’s Hospital Animal Experimentation Ethics Committee (No. 19/07, 19/14), and performed in strict accordance with the National Health and Medical Research Council (NHMRC) of Australia Guidelines on Animal Experimentation. Mice were maintained on a 12-hour light/dark cycle from 6 AM to 6 PM and had unrestricted access to food and water. All efforts were made to minimize animal suffering. We used 3-6-month-old WT male and female animals for our EdU studies.

### PubMed Search

To determine the number of PubMed articles mentioning bromodeoxyuridine (BrdU) and ethynyl deoxyuridine (EdU), a public search was conducted on 13 January 2025 using the PubMed database (https://pubmed.ncbi.nlm.nih.gov/). Search results per year were downloaded as CSV files and subsequently plotted for analysis.

### Reagents

Recombinant human (rh) PDGF-BB (100-14B, PeproTech Inc., US) was reconstituted in PBS1X containing 0.01% BSA, according to the supplier’s instructions, and used as indicated in the corresponding figures. Lyophilized rhFGF-2, research-grade (130-093-837, Miltenyi Biotec) was reconstituted with deionized sterile-filtered water. Other reagents, unless otherwise indicated, were purchased from Sigma-Aldrich.

### Cell Culture and Cell Growth of C3H/10 T1/2 and C2C12 Cells

The murine mesenchymal stromal cell line C3H/10T1/2, Clone 8 (CCL-226, ATCC, VA, USA) and C2C12 myoblast cell line (ATCC CRL-1772) were grown at 37°C in 5% CO□ and cultured in DMEM, high glucose, GlutaMAX™ Supplement (Cat. No. 10566016), supplemented with 10% (v/v) heat-inactivated fetal bovine serum (FBS; Hyclone, UT, United States) and antibiotics (Penicillin-Streptomycin, Cat. No. 15140122, Gibco by Life Technologies). Cells were passaged every other day and used between passages 10-20. Additionally, cells were treated with rhPDGF-BB and rhFGF-2 in DMEM supplemented with 1% FBS and antibiotics at 37°C in 5% CO□, at concentrations and times indicated in the corresponding figure legend. Our cell cultures were periodically tested for mycoplasma contamination using PCR at the Garvan Institute of Medical Research.

### hiPSC Cell Culture

The hiPSC line PB010.5 with the incorporation of a reporter YFPhGEMININ/mCherryhCDT1 FUCCI cassette into the EEF2 locus, which exhibited no karyotypic abnormalities, was generated by the iPSC Derivation & Gene Editing Facility (Murdoch Children’s Research Institute, Melbourne, Australia). These hiPSCs were split using ReLeSR™ (100-0483, STEMCELL Technologies) and maintained in mTeSR™ Plus (100-0276, STEMCELL Technologies) with 0.5% v/v Penicillin-Streptomycin (15140122, Gibco) under feeder-free conditions on hESC pre-screened Matrigel (354277, Corning). hiPSCs were used between passages 30 to 45. To ensure the quality and pluripotency status of hiPSCs over long-term maintenance, we periodically conducted standard flow cytometric immunophenotyping (Miltenyi Biotec-tested panel 25, Miltenyi Biotec), as well as assessed the growth rate and morphology of hiPSC colonies. hiPSCs were periodically tested for mycoplasma contamination using PCR at the Garvan Institute of Medical Research.

### Cancer Cell Growth Protocol and Conditions

Cancer cell lines were grown at 37°C in 5% CO□ and cultured in DMEM, high glucose, GlutaMAX™ Supplement (Cat. No. 10566016), supplemented with 10% (v/v) heat-inactivated fetal bovine serum (FBS; Hyclone, UT, United States) and antibiotics (Penicillin-Streptomycin, Cat. No. 15140122, Gibco by Life Technologies). Panc1 cells (passage 20) were cultured at a density of 16,000 cells/cm², MCF7 cells (passage 60) at 25,000 cells/cm², MDA-MB-231 cells (passage 25) at 12,000 cells/cm², and MIA PaCa2 cells (passage 16) at 16,000 cells/cm². Cells were pre-plated onto 6-well plastic tissue culture dishes and incubated at 37°C in a 5% COC atmosphere for about 24 or 48 hours. Before the EdU thymidine analog treatment, the growth medium was replaced with a fresh medium for two hours. Subsequently, 10 µM of EdU was added to the cultures as indicated in the figure legends and text.

### Fixation Protocol for Flow Cytometry: Cultured Cell Lines and hiPSCs

For fixing cell lines (C2C12, NIH3T3, C3H-10T1/2, cancer cell lines) and hiPSCs for flow cytometry, cells in 6-well plates were processed as follows. The media was removed, and each well was washed once with 1 mL of PBS. Then, 1 mL of TrypLE was added to each well, and the plates were incubated for 5-7 minutes at 37°C. The cells were pipetted up and down a few times per well to obtain a single-cell suspension, which was corroborated using a microscope. Next, 1 mL of cold FACS Buffer (PBS 1×, 2% FBS v/v, 2 mM EDTA pH 7.9) was added to each well to inactivate the TrypLE. The cells from each well were transferred to separate 2 mL tubes and spun at 500g for 5 minutes (200g for 3 minutes for hiPSCs). The supernatant was removed, and the pellets were resuspended by flicking the tubes. Then, 1 mL of FACS Buffer was added to each well. The tubes were checked for clumps by looking at them against a light source, and any clumps present were removed by flicking the tubes. 4% PFA was added to a final concentration of 2% in each tube, and the cells were fixed for 10 minutes at room temperature. After fixation, the cells were washed with abundant PBS, and the tubes were spun at 400g for 4 minutes (200g for 3 minutes for hiPSCs). The supernatant was removed, and the pellets were resuspended by flicking. Finally, 1-2 mL of PBS with 0.02% Sodium Azide was added to each tube, and the samples were stored at 4°C. The volumes were adapted when using smaller well plates.

### Cytospin of Cells in Suspension

After mounting CytoSep™ Cytology Funnels (base holder and fluid chamber) for the Sakura Cyto-Tek® Cytocentrifuge (Model 4323, Sakura) and a white filter for the Sakura cytology funnel (M963FW, Simport), PFA-fixed cells in suspension were centrifuged (2,000 rpm, 3 min) onto Epredia™ SuperFrost Plus™ Adhesion slides (12312148, Fisher Scientific). Following the creation of a hydrophobic barrier with a PAP pen (ab2601, Abcam), the cells were ready for staining. We typically used 0.1 to 0.8 mL of cells in suspension.

### OpenEMMU Click Chemistry for Flow Cytometry Combined with Antibody Staining

For flow cytometric analysis of cells, the required number of fixed cells was transferred into 1.5 mL Eppendorf tubes. We typically used between 100,000 to 1,000,000 cells. Each tube received 1 mL of 1% BSA in PBS, followed by centrifugation at 500g for 5 minutes. The supernatants were discarded, and the pellets were resuspended by flicking. Subsequently, 500 µL of 1X in-house permeabilization/wash buffer (0.2% (w/v) saponin containing either 4% FBS (v/v) or 2% NCS (v/v), 1% (w/v) BSA and 0.02% (v/v) Sodium Azide in PBS) was added to each tube, and the samples were incubated at room temperature (RT) for 10 minutes. After incubation, the tubes were centrifuged again at 500g for 5 minutes. The supernatants were discarded, leaving 50 µL in each tube, and the pellets were resuspended by flicking. The EdU mix was then prepared by sequentially adding the following components: 909 µL PBS, 80 µL of Copper(II) sulfate pentahydrate (209198, Sigma Aldrich; 10 mM fresh or frozen stock, final concentration 800 µM), 1 µL Fluor-Azide (200 µM stock, final concentration 200 nM), and 10 µL freshly made L-Ascorbic acid (100 mg/mL frozen stock, final concentration 5.7 mM). 300-450 µL of the EdU mix was added to each tube, and the samples were incubated at RT for 30 minutes, covered from light. Following this, the tubes were washed twice with 1 mL of 1% BSA in PBS and once with 0.5 mL of 1X permeabilization/wash buffer, and centrifuged at 500g for 5 minutes in between washes. The supernatants were discarded, leaving 50 µL in each tube, and the pellets were resuspended by flicking. 100 µL of primary or conjugated antibodies, prepared in 1X permeabilization/wash buffer, was added to the relevant tubes, and the samples were incubated overnight at 4°C. The next day, the tubes were washed once with 1 mL of 1% BSA in PBS and once with 0.5 mL of 1X perm/wash buffer, followed by centrifugation at 500g for 5 minutes. The supernatants were discarded, leaving 50 µL in each tube, and the pellets were resuspended by flicking. When using secondary antibodies, 50 µL of them, prepared in 1X permeabilization/wash buffer, was added to the relevant tubes, and the samples were incubated at RT for 60-120 minutes, covered. After this incubation, the tubes were washed once with 1 mL of 1% BSA in PBS and once with 0.5 mL of 1X perm/wash buffer. The supernatants were discarded, leaving approximately 100 µL in each tube, and the pellets were resuspended by flicking. A 1:2500 dilution of Vybrant™ DyeCycle™ Violet (V35003, 5 mM) in 1X perm/wash buffer was prepared, and 100 µL was added to each tube for a final concentration of 1:5000. The tubes were covered and taken for analysis by flow cytometry. Details about the AZDye fluorescent probes and antibodies used in this study can be found in the Supplementary Materials (Table S1).

### In-house OpenEMMU Click Chemistry for Immunofluorescence and Antibody Staining

Cells, either in a cytospin slide and/or PhenoPlate 96-well, black, optically clear flat-bottom plates (Revvity), were washed with 1X PBS and then permeabilized in 1× saponin-based permeabilization/wash buffer for 15 min. During this incubation period, the EdU mix was prepared. After 15 minutes, the 1X permeabilization/wash buffer was removed, and the samples were incubated with the EdU mix (same formula as for flow cytometry) at RT for 30 minutes, covered from the light. Following this, the samples were washed three times with 1X PBS. The samples were then incubated overnight at 4°C with primary or conjugated antibodies prepared in 1X permeabilization/wash buffer. The next day, the samples were washed three times with 1X PBS before adding secondary antibodies, also prepared in 1X permeabilization/wash buffer, and incubated at RT for 60-120 minutes, covered. After this incubation, the samples were washed three times with 1X PBS. Finally, a 1:5000 dilution of Vybrant™ DyeCycle™ Violet in 1X permeabilization/wash buffer was prepared, and the samples were incubated with this solution and stored at 4°C until they were ready for imaging. Details about the AZDye fluorescent probes and antibodies used in this study can be found in the Supplementary Materials (Table S1).

### hiPSC-derived Cardiomyocyte In-house Differentiation Protocol

On Day −2, hiPSCs were dissociated using pre-warmed TrypLE (TrypLE Express Enzyme (1×), no phenol red, Cat. No. 12604013, ThermoFisher Scientific, USA) for 7 min and seeded at a density of 200,000 cells/cm² in 24 or 48-well plates using mTeSR medium supplemented with 5 µM Y-27632. On Day −1, the media in the wells was replaced with fresh mTeSR. On Day 0, when the wells were 70-90% confluent, they were washed once with PBS and 6 µM CHIR99021 in RPMI+B27(-ins)+Glutamax was added. Approximately 24 hours later, on Day 1, the wells were washed once with PBS, and RPMI+B27(-ins)+Glutamax with 1 µM CHIR99021 was added. On Day 3/4, the wells were washed once with PBS and RPMI+B27(-ins)+Glutamax supplemented with 200 µM L-Ascorbic acid, 5 µM IWP-2, and 5 µM XAV939 was added. 48 hours after Day 3/4, on Day 5/6, the wells were washed once with PBS and RPMI+B27(-ins)+Glutamax supplemented with 200 µM L-Ascorbic acid added. On Day 7, the media was replaced with RPMI+B27+Glutamax supplemented with 200 µM L-Ascorbic acid. From Day 9 onwards, the media was replaced every 2-3 days with RPMI+B27+Glutamax supplemented with 200 µM L-Ascorbic acid.

### hiPSC-Derived Cardiomyocyte In-house Dissociation Protocol

Cells in 48-well plates were washed with 0.5 mL/well PBS and then incubated with 250 µL/well of room temperature Cardiomyocyte Dissociation Buffer for 30 minutes at 37°C, tapping the plate every 15 minutes. The Cardiomyocyte Dissociation Buffer was composed of 12.5 U/mL Papain (Cat. #10108014001, Sigma Aldrich), 0.5 mM Cysteine, 132.5 U/mL Collagenase II (Cat. #LS004176, Worthington), 0.5% BSA (Cat. #A7906, Sigma), 0.025 mg/mL DNase I (Cat. #10104159001, Sigma Aldrich), and 0.25 U/mL Dispase (Cat. #07913, STEMCELL), all made up in DMEM (+L-Glutamine, +Sodium Pyruvate, +4.5 g/L Glucose). Following this, 500 µL FACS buffer/well was added, and the plate was placed on ice while still in the hood. Cells were gently pipetted up and down 20 times using a P1000 pipette to achieve single-cell status, with an additional 10 pipetting cycles if needed. The cells were then transferred to 15 mL Falcon tubes, pooling wells as required, and excess FACS buffer was added to each tube. The tubes were spun at 300g for 4 minutes, and the supernatant was carefully removed. The cell pellets were resuspended by flicking. Cells were fixed in 2% PFA (prepared by diluting 4% PFA stock in FACS buffer) for 10 minutes. After fixation, excess PBS was added to the tubes, and they were spun down at 300g for 4 minutes. The supernatant was removed, and the pellets were resuspended, ensuring no clumps were present. Finally, sufficient PBS (+0.02% Sodium Azide) was added to each tube, and the cells were stored at 4°C.

### Mouse Dissection for the Spleen, Bone Marrow, and Small Intestine

Mice were asphyxiated in a CO_2_ chamber at a slow fill rate (approximately 20-30% of the chamber volume per minute). Death was confirmed by cervical dislocation and a lack of response to physical contact. After immobilizing the deceased mouse, the chest fur was lightly sprayed with 80% ethanol. The chest cavity was then opened using external dissection forceps and scissors. The spleen and small intestine were removed and transferred to a dish containing ice-cold DMEM (Cat. No. 10566016) for washing. For the small intestine, a 2 cm section of the ileum was cut, and the digested food was removed by flushing the intestine twice with 10 mL of cold PBS1x. The ileum was fixed in 4% PFA for 30 minutes at room temperature, washed with abundant PBS1x, and stored in 100% methanol at −20°C until further FFPE or tissue-clearing processing.

### Splenocytes Isolation, Fixation and Staining

After EdU uptake, spleens were individually washed in cold DMEM (Cat. No. 10566016) and placed in a 100µm cell strainer, then carefully ground until forming a paste-like slurry. Five milliliters of cold FACS buffer were used to wash the slurry as it passed through the strainer. The remaining tiny pieces of spleen were further ground, followed by another 10 mL of cold FACS buffer. The mixture was centrifuged at 400 g for 5 minutes at 4°C, and the supernatant was discarded. The pellet was resuspended by flicking the tube for about 5-7 seconds to ensure complete resuspension. Five milliliters of ACK/RBC lysis buffer was added to the resuspended pellet and incubated for 3 minutes on ice. Ten milliliters of cold FACS buffer were then added to stop the lysis, followed by centrifugation at 400 g for 5 minutes at 4°C, and the supernatant was discarded. The pellet was resuspended by flicking the tube and 5 mL of cold FACS buffer was added. The splenocytes were filtered through a blue cap flow cytometry tube and then fixed in 1% PFA for 10 minutes at room temperature. The cells were washed with FACS buffer and centrifuged at 500 g for 5 minutes, and the supernatant was carefully discarded. The cells were resuspended in 5 mL of 1% BSA in PBS1x and 0.02% (v/v) Sodium Azide. OpenEMMU click chemistry was then performed using 500,000 to 1,000,000 total cells according to the OpenEMMU protocol for flow cytometry. After Fc blocking (anti-Mouse CD16/CD32 antibody), antibodies were prepared in a 1X in-house permeabilization/wash buffer and incubated with cells for 1 hour at room temperature. Details about the AZDye fluorescent Picolyl Azide and antibodies used in this study can be found in the Supplementary Materials (Table S1).

### Bone Marrow Cell Isolation, Fixation and Staining

After EdU uptake, the tibia and femur bones of adult WT mice (both male and female) were separated from the surrounding muscular and connective tissue. Both ends of the isolated long bones were cut using a surgical blade, and the bone marrow cells were flushed out with FACS buffer using a syringe. The cells were collected in 2 mL Eppendorf tubes, filtered through a pluriStrainer Mini 40µm into a new tube, and then fixed in 1% PFA. OpenEMMU click chemistry was then performed using 500,000 to 1,000,000 total cells as described above. After Fc blocking (anti-Mouse CD16/CD32 antibody), antibodies were prepared in a 1X in-house permeabilization/wash buffer and incubated with cells for 1 hour at room temperature. Details about the AZDye fluorescent Picolyl Azide and antibodies used in this study can be found in the Supplementary Materials (Table S1).

### Embryonic Heart Cell Isolation and Fixation

Following embryonic EdU uptake, wild-type (WT) embryonic hearts from both male and female mice were isolated and cleaned of surrounding tissue. The hearts were then cleared of blood using cold PBS. Each heart was dissociated into single cells with the *Cardiomyocyte Dissociation Buffer* (30 minutes at 37°C). The dissociated cells were collected in 2 mL Eppendorf tubes, filtered through a pluriStrainer Mini 100 µm (43-10100-50, pluriSelect) into a new tube, and fixed in 1% PFA for 10 minutes. OpenEMMU click chemistry and antibody staining was subsequently performed on 100,000 to 500,000 total cells as previously described.

### Multi-parametric Flow Cytometry Analyses

Multi-parametric flow cytometry analyses were performed using a BD LSRFortessaTM X-20 Cell Analyzer and a BD FACSymphony A5 High-Parameter Cell Analyzer, both equipped with five excitation lasers (UV 355 nm, Violet 405 nm, Blue 488 nm, Yellow/Green 561 nm, and Red 633 nm). FSC-H versus FSC-A, FSC-H versus FSC-W, and SSC-H versus SSC-W cytograms were used to discriminate and gate out doublets/cell aggregates during sorting or analysis. Data were collected using BD FACSDiva™ Software. For optimal DNA dye signal detection and cell cycle progression analyses, an event concentration of <1,000 events/seconds was used, and 10,000-50,000 events were captured for most cell types. For analysis of complex organs and tissues, including the spleen and bone marrow, around 500,000-1,000,000 events were captured. All flow cytometry data were analyzed using FlowJo Portal (version 10.8.1, BD) using macOS Monterey. Automated or manual compensation was done only when exclusively required.

### Confocal Laser Scanning Microscopy

Confocal laser scanning microscopy of cells and tissues was performed using a Zeiss LSM900 inverted confocal laser scanning microscope, which includes an upright Zeiss Axio Observer 7, Colibri 5 solid-state LED fluorescence light sources (solid-state laser lines: 405, 488, 561, 640), two Gallium Arsenide Phosphide photomultiplier tubes (GaAsP-PMT), and a motorized stage controlled by ZEN blue 3.4 software. Regular or tile images were acquired using objectives ranging from ×10 (0.45 NA with a WD 2.0, Air Plan-APO UV-VIS–NIR), ×20 (0.8 NA with a WD 0.55, Air Plan-APO UV-VIS–NIR), ×40 (1.3 NA with a WD 0.21, Oil Plan-APO DIC-UV-VIS–IR), and ×63 (1.2 NA with a WD 0.19, Oil Plan-APO DIC-UV-VIS–IR). The ×40 and ×63 objectives were used with immersion oil ImmersolTM 518 F (433802-9010-000, Zeiss). For z-stack 3D imaging, z-step sizes ranged from 0.25-1 μm, with images acquired under confocal settings using a motorized focus drive. Laser power during imaging was kept below 3.5%. When indicated, some tissue FFPE sections were imaged using a Leica Thunder Imager and Leica Image Files (LIFs) were imported directly in Fiji.

### Timed Mating and Dissection of the Mouse Embryo for FFPE

Timed mating was performed between wild-type (WT) male and female mice, with pregnancy confirmed by the detection of a copulation plug. Pregnant females received an intraperitoneal injection of 20 mg/Kg EdU in sterile 0.9% NaCl solution between embryonic days E10 and E14.5.

Three hours post-injection, the pregnant mice were sacrificed as previously described, and embryos were collected in glass Petri dishes containing ice-cold PBS1x. The embryos were briefly rinsed in ice-cold PBS1x to remove any residual blood, then immersed in 4% PFA and placed on a roller for 3 hours at room temperature. Embryonic hearts were isolated by surgical removal. The PFA solution was then replaced with PBS1x, and whole embryos or isolated hearts were stored in 100% methanol at −20°C until further FFPE processing.

### Formalin-Fixed Paraffin-Embedded (FFPE) Tissue Processing and Embedding

All tissues, organs, embryos, and specimens designated for FFPE were initially fixed in 4% paraformaldehyde (PFA) and subsequently preserved in 100% methanol for later processing. Tissue samples were then processed using the Leica Peloris 3, following one of the standard protocols, which lasted either 4 or 8 hours and involved the same steps and reagents. The samples progressed through 70% ethanol, 90% ethanol, and 100% ethanol, followed by Xylene at room temperature, and finally paraffin at 60°C. The protocol was chosen based on the sample type and size. Generally, small samples (less than 3mm small/thin) were processed on the 4-hour run. Samples were then embedded using the Leica HistoCore Arcadia H&C.

### Sectioning, H&E, and IHC

Blocks were sectioned at 5µm using a Leica RM2235 microtome. Sections were placed on either plain glass slides or positively charged slides and incubated for 2 hours in a 60°C oven to ensure maximum adhesion. H&E is a regressive stain, and the standard H&E procedures were followed using the Leica ST5010 Autostainer XL with Hematoxylin (Hematoxylin Harris non-toxic (acidified); Australian Biostain) and Eosin (Eosin Phloxine Alcoholic 1%; Australian Biostain).

### Deparaffinization and Antigen Retrieval Protocol for FFPE Tissues

We used an optimized protocol for FFPE, deparaffinization, and antigen retrieval (Zaqout, Becker, and Kaindl, 2020). In brief, FFPE slides were incubated in Xylene for 15 minutes, repeated three times. This was followed by incubation in a 1:1 mixture of Xylene and ethanol for 5 minutes. Slides were then incubated in 100% ethanol for 3 minutes, repeated twice, followed by 95% ethanol for 3 minutes, repeated twice, and 70% ethanol for 3 minutes, repeated twice. Finally, slides were washed in PBS for 3 minutes, and repeated twice. For antigen retrieval, Citrate Buffer was prepared by dissolving 2.94 g of Trisodium Citrate Dihydrate in 1000 mL of distilled water. The pH was adjusted to 6 using 1M HCl, and 0.5 mL of Tween-20 was added. The Citrate Buffer was heated on a hotplate, and when the temperature reached 95°C, slides were placed in the beaker and incubated for 15 minutes. After incubation, slides were removed from the buffer and allowed to cool at room temperature for 10 minutes. The slides were then ready to be stained.

### Ethyl Cinnamate Tissue Clearing and Click Labeling of Mice Intestines

Tissues stored in 100% methanol at −20°C were rehydrated through successive 5-minute incubations in 75%, 50%, and 25% methanol prepared in PBSTx (PBS + 0.3% Triton X-100). This was followed by two washes in PBSTx for 5 minutes each, and then two washes in PBS for 5 minutes each. The tissues were de-pigmented in 3% H_2_O_2_ (prepared in 0.8% KOH) for 5 minutes at room temperature. After de-pigmentation, the tissues were washed four times in PBS. They were then permeabilized in PBSTx for 60 minutes at room temperature and subsequently overnight at 4°C. The next day, most of the supernatant was removed, and 500 µL of fresh OpenEMMU reaction mix (prepared in PBSTx or Saponin-based perm/wash buffer) was added to each tube using AZDye 488. The tubes were incubated and covered at room temperature for 60 minutes. Following this, the tissues were washed four times in PBS. Five hundred microliters of a solution containing conjugated antibodies or CF®640R WGA (29026, Biotium) (prepared in Zebrafish permeabilization/wash buffer: 0.4% Saponin, 4% FBS, 1% BSA, 10% DMSO, 1% Triton X-100, 0.02% Sodium Azide in PBS) was added to each tube and incubated covered at room temperature for 30 minutes. The tissues were then washed four times in PBS. The tubes were dehydrated through successive 30-minute incubations at room temperature with rocking in 30%, 50%, and 70% propane-2-ol (prepared in PBS), followed by 100% propane-2-ol. The tubes were then dehydrated overnight in 100% propan-2-ol at 4°C. The next day, propan-2-ol was replaced with ethyl cinnamate (112372, Sigma-Alrich), and the tubes were incubated and covered on a rocker. The ethyl cinnamate was replaced after 1 hour to remove any traces of propan-2-ol. The tubes were then allowed to incubate at room temperature, and covered on a rocker overnight. Finally, the intestines were imaged using laser confocal imaging.

### Deep Clearing and Click Labeling of Mice Embryos

Embryos stored in methanol were rehydrated through consecutive 10-minute incubations in 75%, 50%, and 25% methanol in PBSTx0.3 (0.3% Triton X-100 in PBS). This was followed by two washes in PBSTx0.3. The embryos were then permeabilized in ice-cold acetone overnight at −20°C. After permeabilization, embryos were washed four times in PBS for 5 minutes each. Embryos were transferred into glass vials and 5 mL of Solution-1 (25% Urea, 9% THEED, 5% Triton X-100 in water) was added. The vials were incubated on a rocking platform with gentle rocking at 37°C for 5 days, with fresh Solution-1 added every 2-3 days or whenever the solution appeared yellowish. Following this, embryos were washed three times with 4 mL of PBS for 10 minutes each. Blocking was performed by adding 4 mL of blocking PBSTx (4% FBS, 1% Triton X-100, 1% BSA in PBS) to each vial and incubating on a rocking platform with gentle rocking at 37°C overnight. Embryos were then transferred into 2 mL Eppendorf tubes and washed twice in PBSTx0.3 for 10 minutes each. An EdU mix was prepared using PBSTx1 (1% Triton X-100 in PBS) and 1 mL was added to each tube. The embryos were incubated on a rocking platform with gentle rocking at 37°C overnight. Following EdU incubation, tubes were washed four times in PBSTx0.3 for 15 minutes each on a rocking platform with gentle rocking at 37°C. Primary antibodies were prepared in Zebrafish perm/wash buffer (0.4% Saponin, 4% FBS, 1% BSA, 10% DMSO, 1% Triton X-100, 0.02% Sodium Azide in PBS). 500 µL of the antibody mix was added to each tube and incubated on a rocking platform with gentle rocking at 37°C for 7 days. This was followed by four washes in PBSTx0.3 for 15 minutes each on a rocking platform with gentle rocking at 37°C. Secondary antibodies, including Vybrant Violet, were prepared in Zebrafish perm/wash buffer. 500 µL of the secondary antibody mix was added to each tube and incubated on a rocking platform with gentle rocking at 37°C for 7 days. The tubes were then washed four times in PBS for 1 hour each and then overnight on a rocking platform with gentle rocking at 37°C. Embryos were RI matched by incubating them in a solution of 7M Urea in PBS on a rocking platform with gentle rocking at 37°C for 5 days. Finally, the objects were analyzed using Lightsheet Microscopy.

### Deep Clearing and OpenEMMU Labeling of Mouse Embryonic Hearts

Hearts stored in methanol were rehydrated through consecutive 5-minute incubations in 75%, 50%, and 25% methanol in PBSTx0.3 (0.3% Triton X-100 in PBS). This was followed by two washes in PBSTx0.3 for 5 minutes each. The hearts were then permeabilized in ice-cold acetone overnight at - 20°C. After permeabilization, the hearts were washed four times in PBS for 5 minutes each. The hearts were transferred into Eppendorf tubes, and 1 mL of Solution-1 (25% Urea, 9% THEED, 5% Triton X-100 in water) was added to each tube. The tubes were incubated on a rocking platform with gentle rocking at 37°C overnight. Following this, the hearts were washed three times with 1 mL of PBS per tube for 5 minutes each. Blocking was performed by adding 1 mL of blocking PBSTx (4% FBS, 1% Triton X-100, 1% BSA in PBS) to each tube and incubating on a rocking platform with gentle rocking at 37°C overnight. The hearts were then washed twice in PBSTx0.3 for 5 minutes each. OpenEMMU mix was prepared using PBSTx1 (1% Triton X-100 in PBS), and 600 µL was added to each tube. The hearts were incubated and covered on a rocking platform with gentle rocking at 37°C overnight. Following incubation, the tubes were washed four times in PBS for 5 minutes each, covered on a rocking platform with gentle rocking at 37°C. Primary antibodies were prepared in Zebrafish perm/wash buffer. 200 µL of the antibody mix was added to each tube and incubated covered on a rocking platform with gentle rocking at 37°C over the weekend (approximately 3 days). This was followed by three washes in PBSTx0.3 for 10 minutes each on a rocking platform with gentle rocking at 37°C. 500 µL of the secondary antibody mix (prepared in Zebrafish perm/wash buffer) was added to each tube and incubated covered on a rocking platform with gentle rocking at 37°C overnight. The tubes were then washed three times in PBSTx0.3 for 30 minutes each on a rocking platform with gentle rocking at 37°C. The hearts were RI matched by incubating them in a solution of 7M Urea in PBS, covered on a rocking platform with gentle rocking at 37°C for 30 minutes. Finally, the hearts were analyzed using light-sheet microscopy.

### Deep Clearing and Click Labeling of Mice Limbs

Limbs were rehydrated by incubating in successive dilutions of methanol (75%, 50%, and 25%) prepared in PBSTx0.3 (0.3% Triton X-100 in PBS) for 10 minutes each. This was followed by three washes in PBSTx for 5 minutes each. The limbs were then permeabilized in ice-cold acetone overnight at −20°C. After permeabilization, the limbs were washed four times in PBS. The limbs were incubated in Solution-1 (25% Urea, 9% THEED, 5% Triton X-100 in water) for 6 hours at 37°C on a rocker. Following this, the limbs were washed four times in PBS. They were then permeabilized overnight in PBSTx1 (1% Triton X-100 in PBS) at room temperature on a rocker. Most of the PBSTx was removed, and 500 µL of the OpenEMMU reaction mix (prepared using PBSTx1) was added to each tube. The tubes were incubated at room temperature on a rocker for 120 minutes. After the reaction, the limbs were washed four times in PBS. The limbs were then incubated overnight in conjugated antibodies and DNA dyes prepared in Zebrafish perm/wash buffer at room temperature on a rocker. After incubation, the limbs were washed four times in PBS. Finally, the samples were RI matched by incubating them in a solution of 7M Urea in PBS at room temperature on a rocker for an appropriate amount of time and imaged using Light Sheet fluorescence microscopy.

### Light Sheet-based Fluorescence Microscopy (LSFM) for Imaging of Large Specimens

All LSFM-dedicated samples were mounted in a 1-2% low melting agarose solution (16520050, ThermoFisher) in PBS by directly mixing the object with the agarose, then pumping the mixture into sample embedding glass cylinder capillaries, selected based on the object’s size and volume (inner diameter of capillary: size 1/∼0.68 mm, size 2/∼1 mm, size 3/∼1.5 mm, size 4/∼2.15 mm, Zeiss). After positioning the specimens using a camera with LED illumination, three-dimensional images were acquired on a LightSheet Z.1 (Zeiss). The system was equipped with multiple solid-state laser lines (405 nm, 445 nm, 488 nm, 515 nm, 561 nm, 638 nm) and two 16-bit sCMOS PCO.Edge cameras for simultaneous dual-color acquisition (1920 x 1920 pixels). The light sheet was focused onto the specimen using Illumination Optics Lightsheet Z.1 5x/0.1 on both the left and right sides. Images were captured with Lightsheet Z.1 detection optics 5x/0.16 NA EC Plan NEOFLUAR 1.33/1.45 adaptive rings, with a z-step of 8 to 11 μm, a pixel size of 6.5 nm, and a Filter Module LBF 405/488/561/638. Dual-side fusion and relevant processing were performed offline using ZEISS ZEN Black software for Z.1. All images were further post-processed, handled, and automated 3D multiview reconstructed and merged using Fiji software (ImageJ2, version: 2.14.0/1.54f).

### Generation of iPSC-derived Cardiac Organoids

Human induced pluripotent stem cells (hiPSCs) specifically used in human cardiac organoids were cultured under feeder-free conditions. Briefly, two hiPSC lines (CMRIi0013-A-6 9CMRI, Stem Cell and Organoid Facility and HPS10314I-hoik_1 (ECCA)) were maintained on E8 media (Thermofisher, #A1517001) and Geltrex matrix (ThermoFisher, #A1413302) and passaged as clumps with ReleSR (Stem Cell Technologies, #100-0483) upon reaching 60-70% confluence. Cardiac organoids were generated from 70% confluent hiPSCs cultures according to the protocol described by Hofbauer et al (2021) with some modifications. Briefly, hiPSCs were dissociated using Tryple Express (Thermofisher, #12604021) to single cells. Ten thousand cells per 200µL of E8 (Thermofisher, #A1517001) supplemented with 10µM Y-27632 ROCK inhibitor (Stem cell Technologies, #72304) were seeded on a low-binding 96-well U-bottom plate (Thermofisher, #174925). The plate was then centrifuged twice (clockwise and anti-clockwise) at 200g for 5 minutes. At day 1 (D1) of differentiation, mesoderm induction was initiated. E8 media was changed to chemically defined media (CDM) (IMDM + GlutaMAX (Thermofisher, #31980030) and Ham F12 + GlutaMAX (Thermofisher, #11765054) in a 1:1 ratio supplemented with 5mg/mL BSA (Merck, #A1470), 1X Lipid Mix (Merck, #L5146), 1X ITS (Thermofisher, #41400045), 95U/mL PenStrep (Thermofisher, #15070063), 0.4µg/mL Amphotericin (Thermofisher, #15290018) and 450µM 1-Monothioglycerol (Merck, #M6145)) supplemented with 4µM and 6µM CHIR99201 (Stem cell Technologies, #72054) for HPS10314I-hoik_1 and CMRIi0013-A-6 respectively, 50ng/mL Activin A (R&D Systems, #338-AC-10), 10ng/mL BMP4 (R&D Systems, #314-BP-010), 30ng/mL FGF2 (R&D Systems, #233-FB-025). Three days later (D3), cardiac mesoderm induction started by adding fresh CDM supplemented with 10 ng/mL BMP4 (R&D Systems, #314-BP-010), 8ng/mL FGF2 (R&D Systems, #233-FB-025), 5µM IWP-2 (Tocris, #353-310), 0.5µM Retinoic Acid (Merck, #R2625, 10µg/mL Insulin (Merck, #I9278), 200 ng/mL VEGF (Peprotech, #AF-100-20). Daily media changes from D3-D7 were performed. On D7, media was changed to CDM supplemented with 10 ng/mL BMP4 (R&D Systems, #314-BP-010), 8ng/mL FGF2 (R&D Systems, #233-FB-025), 10µg/mL Insulin (Merck, #I9278) and 100 ng/mL VEGF (Peprotech, #AF-100-20). No media change was required until D10. From D10 onwards, cardiac organoids were maintained on CDM supplemented with 10µg/mL Insulin (Merck, #I9278) and 100 ng/mL VEGF (Peprotech, #AF-100-20), and feeding was performed every other day. By D12, beating cardiac organoids were transferred to a 6-well plate pre-treated with anti-adherence rinsing solution (Stem cell Technologies, #07010) and placed on a shaker at 80 RPM. Samples were collected on D12 and D19 for EdU labeling and analysis. hCOs were labeled with 10µM EdU for 24 hours, washed with PBS, and fixed in 4% PFA for 30 minutes at room temperature. Subsequently, hCOs were stored in 100% methanol at −20°C.

### Tissue Clearing and Click Labeling of 3D Heart Organoids

hCOs were rehydrated by incubating in successive dilutions of methanol (75%, 50%, and 25%) prepared in PBSTx (0.3% Triton X-100 in PBS) for 5 minutes each. This was followed by three washes in PBSTx for 5 minutes each. Organoids were then permeabilized in ice-cold acetone overnight at −20°C. After permeabilization, organoids were washed three times in PBS for 5 minutes each. Organoids were then incubated in Solution-1 (25% Urea, 9% THEED, 5% Triton X-100 in water) (DEEP-Clear; Pende et al., 2020) overnight (or over the weekend) at 37°C with gentle rocking. To pellet the organoids, tubes were spun briefly in a tabletop microfuge. One milliliter of the supernatant was removed and replaced with PBS. After a few minutes, the organoids became more visible as they turned opaque. Organoids were washed three more times with PBS for 5 minutes each. Blocking was performed by adding 1 mL of blocking PBSTx (4% FBS, 1% BSA, 1% Triton X-100 in PBS, filtered) to each tube and incubating at 37°C on a rocker for 90-120 minutes. Most of the supernatant was then removed. OpenEMMU mix was prepared using 1% Triton X-100 in PBS, and 500 µL was added to each tube. Organoids were incubated at 37°C on a rocker for 3 hours. Following incubation, tubes were washed three times with PBS and once with 1X Zebrafish perm/wash buffer (0.4% Saponin, 4% FBS, 1% BSA, 10% DMSO, 1% Triton X-100, and 0.02% NaN3) for 10 minutes each. All but 100 µL of the supernatant was removed. Primary antibodies were prepared in 1X Zebrafish perm/wash buffer, and 100 µL was added to each tube. Tubes were incubated at 37°C on a rocker for 24-48 hours (or over the weekend). Tubes were then washed four times with PBSTx (0.3% Triton X-100) for 30-60 minutes each (or overnight) at 37°C on a rocker, with the first wash lasting 5 minutes. This was followed by a wash with 1 mL of Zebrafish perm/wash buffer for 10 minutes at 37°C on a rocker. Most of the supernatant was removed. Secondary and/or fluorochrome-conjugated antibodies (plus Vybrant Violet) were prepared in Zebrafish perm/wash buffer, and 200 µL was added to each tube. Tubes were incubated at 37°C on a rocker for 24-48 hours (or over the weekend). Finally, tubes were washed four times with PBSTx for 30 minutes each at 37°C on a rocker, with the first wash lasting 5 minutes.

### Proliferation of human T cells in vitro

#### Subjects and Ethics

Recruitment of healthy adult donors through St Vincent’s Hospital was approved by St Vincent’s Hospital Human Research Ethics Committee (HREC/13/SVH/145 and HREC/10/SVH/130). All participants gave written informed consent.

### Antigens

Polyclonal T cell activator anti-CD3/anti-CD28/anti-CD2 was used as a positive control (1/100 dilution; StemCell Technologies, Vancouver, Canada). Recall antigens for memory CD4 T cells included: influenza vaccine (1/100 dilution; Influvac Tetra, 2018 formulation, Mylan Health, Sydney, Australia); *M. avium intracellulare* lysate (5 µg/ml, CSL, Melbourne, Australia); and trimeric recombinant SARS-CoV-2 spike protein (5 µg/ml; which was produced from a plasmid encoding the spike protein with C-terminal trimerization domain and His tag (gift from the Krammer lab, BEI Resources, NIAID, NIH), transfected into Expi93 cells and protein expressed for 3 days, initially purified using the His tag and Talon resin (ThermoFisher), and further purified on a Superose 6 gel filtration column (GE Healthcare) using an AKTA Pure FPLC instrument (GE Healthcare) to isolate the trimeric protein and remove S2 pre-fusion protein, as previously described (Rouet et al., 2021; Phetsouphanh et al., 2022).

### CellTrace™ Violet and T-cell proliferation assay

Peripheral blood mononuclear cells (PBMC) were isolated from Na Heparin anti-coagulated blood using Ficoll-Hypaque gradient centrifugation, resuspended at a concentration of 10,000,000 cells/ml in PBS, and incubated with CellTrace™ Violet dye (C34571, Thermofisher) at 5 µM for 20 min at RT, according to the manufacturer’s directions, as previously described (Phetsouphanh et al., 2022). Cells were washed once with 5x volume of IMDM/10% human serum. Antigen-specific CD4 T cells proliferating in response to recall antigens were measured in cultures of 300,000 PBMC in 200 µl/well of a 96-well plate, in Iscove’s Modified Dulbecco’s Medium (IMDM 12440053; Thermofisher) containing 10% human serum (kind gift, Dr. Wayne Dyer, Australian Red Cross Lifeblood, Sydney, Australia), incubated for 7 days in a 5% CO_2_ incubator (Phetsouphanh et al., 2022). Different wells contained different antigens as indicated: (i) culture medium only (negative control well); (ii) anti-CD3/anti-CD28/anti-CD2 T cell activator (positive control well); (iii) influenza vaccine (1/100 dilution); and (iv) *M. avium intracellulare* lysate (5 µg/ml); and (v) trimeric recombinant SARS-CoV-2 spike protein (5 µg/ml). After 6 days, cells from the respective cultures were stained with CD3-BV786, CD4-BUV395, CD8-BUV805 and CD25-APC (BD Biosciences), washed once with PBS, fixed in 1% PFA for 30 min at RT and stored at 4°C for optimized OpenEMMU staining with AZDye 488 Picolyl Azide, and acquired on a 5-laser Fortessa X20 (Phetsouphanh et al., 2022). Antigen-specific CD4 T cells were gated as CD3+CD4+CD25^high^CTV^dim^ as previously described (Zaunders et al., 2019). Cultures were classified as positive for antigen-specific CD4 T cells if the proportion of CD25^high^CTV^dim^ % from CD4+ CD3+ T cells was ≥ 1% (Phetsouphanh et al., 2022; Zaunders et al., 2019).

### Zebrafish Maintenance and Breeding

Wild type *Danio rerio* zebrafish (Ekkwill (EK) strain) were maintained at the Victor Chang Cardiac Research Institute. All procedures were approved by the Garvan Institute of Medical Research/St Vincent’s Hospital Animal Ethics Committee under Animal Research Authorities (AEC: 22_25). Adult zebrafish were housed in 3.5 L tanks Z-Hub system (Aquatic Habitats), with a maximum of 30 fish per tank. They were kept in recirculating chlorine-free water at 27 ± 1 °C, exposed to a 14.5:9.5-hour light-dark cycle, and fed twice daily. Fertilized embryos were incubated at 28°C (Memmert, Büchenbach, Germany) in Embryo Media, which consisted of 0.03% (w/v) ocean salt (Aquasonic, Wauchope, Australia), 0.0075% (w/v) calcium sulfate, and 0.00002% (w/v) methylene blue in water, at a density of 60-100 embryos per 25 mL dish. At 5 days post-fertilization (dpf), viable free-swimming zebrafish larvae were visually screened. For EdU experiments, adult zebrafish were intraperitoneally injected with 50 μL of 8 mM EdU once daily at 5, 6, and 7 days post-injury.

### EdU Labeling of Zebrafish Larvae

5dpf zebrafish larvae were incubated in 0.5 mM EdU (with a final DMSO concentration of 1%) for 2 hours at 27°C in zebrafish growth media (E3). Following incubation, the zebrafish were euthanized in 0.4% Tricaine for 5 minutes. They were then washed with PBS and fixed for 1 hour at room temperature in MEMFA (0.1M MOPS, 2mM EGTA, 1mM MgSO4, 3.7% PFA, pH 7.4) or 4% PFA, obtaining similar results. After fixation, the zebrafish were washed twice with PBSTx (PBS + 0.3% Triton X-100) followed by a wash with methanol. The methanol was removed and replaced with fresh methanol. The samples were stored at −20°C overnight or until further processing. They can be stored this way for months.

### Deep Clearing and Zebrafish Click Labeling

5dpf zebrafish larvae were rehydrated by incubating in successive dilutions of methanol (75%, 50%, and 25%) prepared in PBSTx (0.3% Triton X-100 in PBS) for 5 minutes each. This was followed by two washes in PBSTx for 5 minutes each. Zebrafish were then permeabilized in ice-cold acetone overnight at −20°C. After permeabilization, zebrafish were washed 4-5 times in PBS. Zebrafish were then de-pigmented in 3% H_2_O_2_ (prepared in 0.8% KOH) for 6 minutes under a bright desk lamp. Following de-pigmentation, zebrafish were washed 4-5 times in PBS. Zebrafish were then incubated in Solution-1.1 (9% THEED, 5% Triton X-100, 5% urea in distilled water) for 10 minutes at room temperature. After incubation, zebrafish were washed 4-5 times with PBS. Zebrafish were then incubated in zebrafish permeabilization/wash solution (0.4% Saponin, 4% FBS, 10% DMSO, 1% BSA, and 0.02% NaN3 in PBS) for 90 minutes. Most of the perm/wash buffer was removed, and 400 µL of the OpenEMMU click reaction mix (using 1% Triton X-100 in PBS) was added to each tube. Zebrafish were incubated at room temperature for 60 minutes. Following click incubation, zebrafish were washed once in zebrafish permeabilization/wash solution without DMSO, and most of the supernatant was removed. DNA dyes were prepared in zebrafish perm/wash solution without DMSO and added to the tubes. Finally, zebrafish were imaged using a suitable microscope. Fig. 8D, S.Fig. 10 and S.Fig. 6D images were obtained using a Zeiss Axio Vert.A1 FL-LED inverted microscope with a Microscopy Camera Axiocam 305 color (D) Cooled 5 Mpx 2/3" CMOS and objective LD A-Plan 5x/0.15 Ph1 M27 (WD=11.7mm at D=1mm polystyrene).

### Cardiac Resection in Adult Zebrafish

Cardiac resection and EdU uptake protocols were conducted following previously described methods (Ogawa et al., 2021; Sheng, Zheng, Kikuchi, 2021). For EdU fluorescent assessment, fixed and frozen hearts were sectioned sagittally at a thickness of 7 µm using a Leica CM1950 clinical cryostat (Leica Biosystems) and collected onto adhesive slides (472042491, Trajan Scientific and Medical). OpenEMMU was then applied to the slides as previously described above for *In-house OpenEMMU click chemistry for immunofluorescence and antibody staining*, and the nuclei were counterstained with TO-PRO™-3 Iodide (642/661) (T3605, ThermoFisher).

### Statistical Analysis

For statistical analysis, unless otherwise specified, all results obtained from independent experiments are reported as means ± standard errors of means (SEM) of multiple replicates. Unless otherwise indicated, “*N*” in Figure Legends represents the number of animals or independent biological samples or replicates per group.

## AUTHOR CONTRIBUTIONS

Conceptualization: O.C.

Methodology: O.C., C.T., J.Z., N.J.M., I.A.M., A.G.C.

Validation: O.C., C.T., J.Z., N.J.M., I.A.M., A.G.C., R.P.H.

Formal analysis: O.C, C.T.

Investigation: O.C., C.T., J.Z., N.J.M., I.A.M.

Resources: O.C., J.Z., A.G.C., R.P.H.

Supervision: O.C., R.P.H.

Data curation: O.C., C.T., J.Z.

Project administration: O.C., R.P.H.

Funding: O.C., R.P.H.

Writing – original draft: O.C.

Writing – review & editing: All authors.

## Competing interests

The authors declare that they have no competing interests.

**Correspondence** and requests for materials should be addressed to Osvaldo Contreras.

## Acknowledgments

We are grateful to Sarah Hancock and Nigel Turner for providing the Panc1, MIA PaCa-2, MCF7, and MDA-MB-231 cancer cell lines, and to Dawei Zheng and Emily Wong for supplying wild-type 5dpf zebrafish larvae. We also thank Kazu Kikuchi for the slides of injured zebrafish hearts and Bernice Stewart for administrative support. Special thanks to Michael Lovelace for his advice on Thunder microscopy and his enthusiastic support of this work, and to the members of the Harvey Lab for their feedback. We appreciate Cecelia Jenkin and Aaron Hay for their zebrafish husbandry, the VCCRI BioCORE facility members, Scott Page for his guidance on the Lightsheet Z.1, and the Victor Chang Cardiac Research Institute Innovation Centre (funded by the New South Wales Government Ministry of Health). We also thank the Garvan Weizmann Center for Cellular Genomics (GWCCG), the Core Facility Flow and Histopathology Facilities at the Garvan Institute of Medical Research (GIMR) for their infrastructure support, particularly Eric Lam, Anaiis Zaratzian, and Andrew Da Silva. Most figures were created using Adobe Illustrator and Adobe Photoshop 2024, Adobe. Supplementary Figures S2 and S3 were created with BioRender.com.

## Data and materials availability

All data needed to evaluate the conclusions in the paper are present in the paper and/or the Supplementary Materials.

## Supplementary figure legends

**Fig. S1. Reagent optimization for the application of OpenEMMU in DNA replication analysis.** (A) Flow cytometry data displaying the proportion of EdU-labeled C2C12 cells following a 2-hour EdU pulse, assessed across various CuSO_4_ concentrations in the click reaction. (B) Comparative analysis of EdU and DNA fluorescence intensity at different CuSO_4_ concentrations, corresponding to data from panel (A). (C) Flow cytometry histograms illustrating EdU and DNA fluorescence intensities. (D) Flow cytometry results showing the proportion of EdU-labeled hiPSCs after a 1-hour pulse with varying concentrations of L-ascorbic acid (L-Asc) in the click reaction, showing the selected optimal concentration of 1 mg/mL. (E) Analysis of EdU-labeled hiPSCs post a 1-hour pulse using different concentrations of picolyl azide in the click reaction, with the selected concentration of 0.1 µM noted in the text. (F) Flow cytometry data comparing EdU and DNA fluorescence intensity in the presence of 2% NCS or 4% FBS, showing no differences in staining or the proportion of EdU-labeled cells. (G) Flow cytometry analysis of EdU-labeled C2C12 cells, evaluating DNA fluorescence intensity with various DNA dyes and concentrations.

**Fig. S2. Detailed protocol for OpenEMMU click-based detection of DNA-replicating cells via flow cytometry.** Step-by-step guide for performing flow cytometry to detect DNA-replicating cells labeled with EdU using OpenEMMU. The protocol outlines 14 defined steps, designed to simplify implementation and ensure reproducibility in diverse research laboratory settings.

**Fig. S3. Detailed protocol for OpenEMMU click-based detection of DNA-replicating cells via imaging in cell culture plates and slides.** (A) Step-by-step protocol for imaging DNA-replicating cells labeled with EdU using OpenEMMU. The guide provides 11 defined steps to facilitate straightforward implementation and ensure reproducibility across research laboratories. (B) Confocal imaging of EdU-labelled C2C12 myoblasts and C3H10T1/2 mesenchymal cells following a 2-hour EdU uptake in growth media. Nuclei were also stained, and segmentation was performed using the *StarDist* plugin in Fiji, showcasing OpenEMMU’s applicability in high-throughput imaging workflows for analyzing DNA replication and cell proliferation.

**Fig. S4. Application of OpenEMMU to normal and cancer cell lines for analysis of DNA replication.** (A) Confocal microscopy images of EdU+ hiPSCs after a 10-minute EdU pulse, with NUCLEOLIN-mediated nucleolar immunolabeling and DNA staining. The right panels show magnified cells at different S-phase stages. (B) Flow cytometry results show the proportion of EdU-labeled cancer cells after a 2-hour pulse. (C) Relationship between EdU fluorescence intensity and DNA content of different cancer cell types. (D) Flow cytometry results showing p-HH3^S10^ and γH2A.X^S139^ labeling after a 2-hour EdU pulse in which EdU was also co-stained. (E) Confocal microscopy images showing a high proportion of hiPSCs in S-phase after a 2-hour EdU pulse, with OCT3/4 immunolabeling and DNA staining. (F) Flow cytometry analysis of EdU labelling combined with DNA fluorescence in control cells and cells treated with PDGF-BB or FGF-2 for 24 hours.

**Fig. S5. OpenEMMU outperforms commercial kits in evaluating DNA replication and S-phase of the cell cycle.** (A) Flow cytometry results comparing OpenEMMU and the Invitrogen EdU kit in terms of EdU fluorescence intensity and the proportion of EdU-labeled cells after a 2-hour pulse in C2C12 myoblasts. (B) Flow cytometry results comparing EdU fluorescence intensity and the proportion of EdU+ cells in spleenocytes.

**Fig. S6. OpenEMMU for measuring DNA replication in adult organs and tissues in mice.** (A) Gating strategy employed to detect DNA replication in isolated bone marrow cells (top panels) and spleenocytes (bottom panels). The lines indicate the selected gate. (B) Hematoxylin and eosin staining of an ileum paraffin section of the small intestine. (C) OpenEMMU is compatible with ethyl cinnamate (ECi) tissue clearing and also saponin or Triton X-100 permeabilization and 3D confocal laser imaging. EdU and WGA stainings are shown. Several highly proliferating cell regions are shown, including the adult stem cell crypt and the transient amplifying cells within the villi. (D) *Z*-stack confocal laser imaging of EdU signals (color-coded depth projection) in DNA-replicating cells, along with WGA labeling to highlight the smooth-muscle cell layer, which is rich in blood vessels. (E) Multiplexed imaging (cycle 3) of the small intestine related to Fig. 4C.

**Fig. S7. Whole-embryo and organ-specific analysis of DNA replication and mitosis in developing mouse embryos and organs.** (A) FFPE-processed embryonic day 14 mouse embryo and imaged with Thunder tile imaging of EdU signal in DNA replicating cells multiplexed with Ki67, αSMA-Cy3, p-HH3^S10^, and DNA labeling. αSMA immunolabelling highlights the developing musculature, heart cardiomyocytes, and smooth muscle cells across multiple organs and tissues. (C) FFPE-processed embryonic day 11.5 mouse embryo imaged with confocal tile imaging of EdU signal in DNA replicating cells, multiplexed with Ki67, αSMA-Cy3, and DNA labeling. (C) Individual channels of each staining are shown in panel B. (D) FFPE section of an E14.5 heart, visualized using tiling confocal microscopy. The section underwent four rounds of immunostaining (4-plex, cycle 3; and 4-plex, cycle 4), with OpenEMMU applied during the second cycle, followed by IBEX starting from the second cycle onwards.

**Fig. S8. Application of OpenEMMU to embryonic heart cells and 3D imaging of forelimb development.** (A) Diagram of an embryonic day 14 (E14) mouse heart, illustrating the steps involved in the OpenEMMU protocol for single-cell imaging. (B) Confocal microscopy images of EdU+ cardiac cells following a 3-hour EdU pulse, with WGA labeling and DNA staining. (C) Proportion of EdU-labelled cells in an E14 mouse heart (n=4). (D) Confocal microscopy images of EdU-labelled cardiomyocytes (cTnT+) after a 3-hour EdU pulse, with DNA staining. The arrows show a pair of EdU+ cardiomyocytes. (E) Light sheet fluorescence imaging of EdU multiplexed with αSMA-Cy3 and p-HH3^S10^ immunolabelling, along with DNA staining of an E14 forelimb, shown from a dorsal view (same as Fig. 6D). The right panels show magnified images of each separated staining, demonstrating the high resolution of EdU labeling in DNA-replicating cells as well as immunolabelling. (F) *YZ*-axis light sheet fluorescence imaging showing that tissue clearing combined with OpenEMMU enables deep imaging, allowing visualization of extensor myofibers, DNA-replicating cells, and mitotic cells.

**Fig. S9. DNA replication in self-organizing 3D human cardiac organoids.** (A) FFPE-processed day 12 hCO and confocal tile imaging of EdU fluorescence in DNA replicating cells. NKX2-5 and cTnT expressing cardiomyocytes are highlighted in the boxed area. NKX2-5/EdU double-positive cells are also shown (magnified box). A tube-like cardiac chamber is also shown (bottom panels).

**Fig. S10. DNA replication in zebrafish larvae and under apical transection of an adult zebrafish heart.** (A) Immersion protocol aimed at labeling DNA synthesis with EdU for 2h in 5dpf developing zebrafish larvae, and processing using FFPE. (B) FFPE-processed larva sagittal cross section and light imaging of H&E staining. (C) FFPE-processed larva sagittal cross section (same as Fig. 8B) and tile imaging with Thunder Leica computational clearing of EdU signal in DNA replicating cells. αSMA immunolabeling depicts the developing musculature and growing myotomes. Nuclei were stained for DNA. (D) Whole-mount micrographs showing a growing Danio rerio 5dpf larva with or without DEEP-Clearing. (E, F) LSFM of a whole 5dpf zebrafish larva labeled with EdU and DNA dye, showing a lateral (E) and ventral view (F). (G) EdU, cTnT-AF647, and DNA labeling revealing proliferative cells after 7 days of apical ventricular resection in an adult zebrafish heart, visualized by confocal microscopy. The dashed lines surround the regenerating apical area of the ventricle.

