## Supplemental Figures for "OpenEMMU: a versatile, open-source EdU multiplexing methodology for studying DNA replication and cell cycle dynamics"

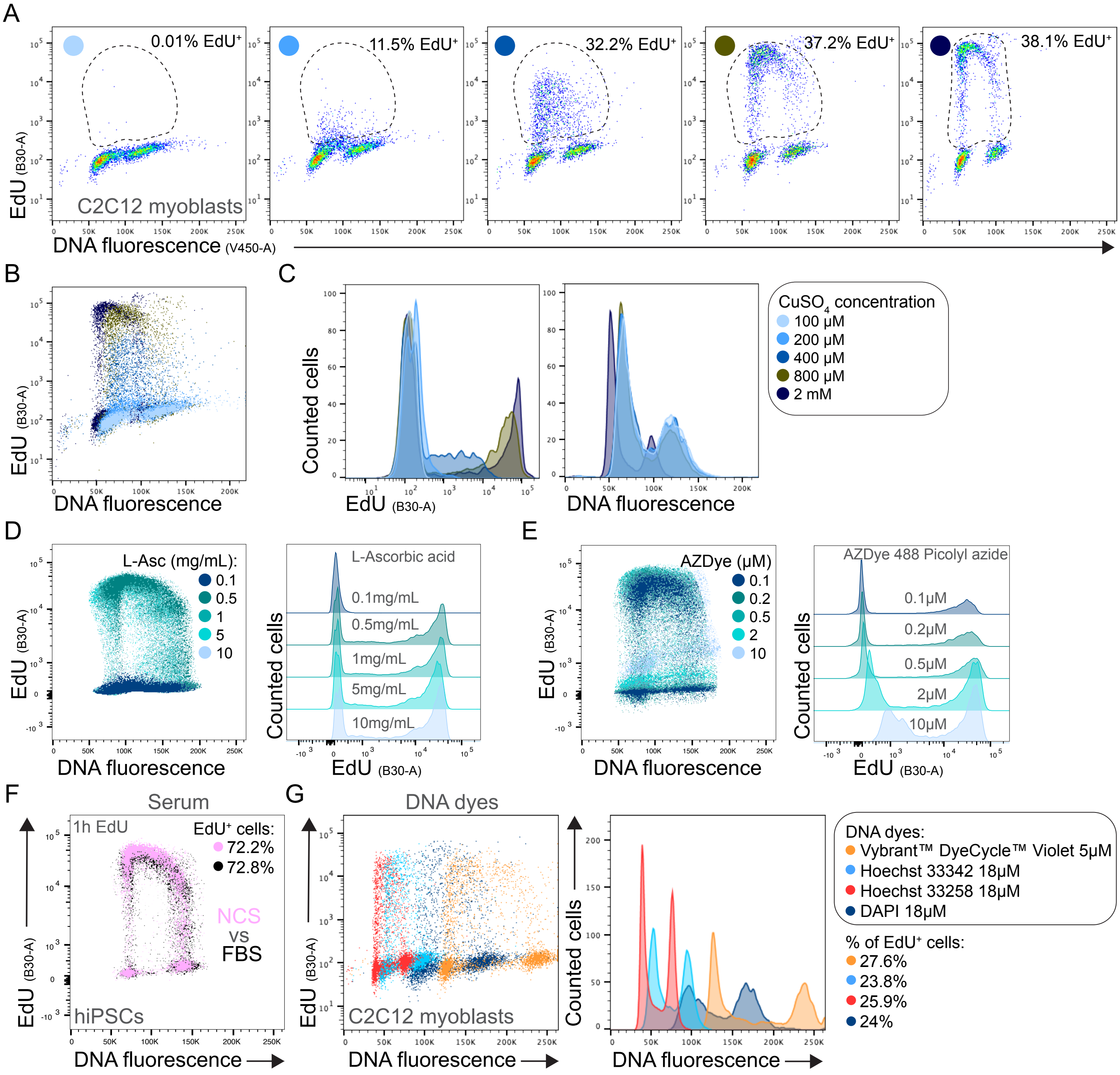


**Fig. S1. Reagent optimization for the application of OpenEMMU in DNA replication analysis.** (A) Flow cytometry data displaying the proportion of EdU-labeled C2C12 cells following a 2-hour pulse, assessed across various CuSO4 concentrations in the click reaction. (B) Comparative analysis of EdU and DNA fluorescence intensity at different CuSO4 concentrations, corresponding to data from panel (A). (C) Flow cytometry histograms illustrating EdU and DNA fluorescence intensities. (D) Flow cytometry results showing the proportion of EdU-labeled hiPSCs after a 1-hour pulse with varying concentrations of L-ascorbic acid (L-Asc) in the click reaction, highlighting the selected optimal concentration of 1 mg/mL. (E) Analysis of EdU-labeled hiPSCs post a 1-hour pulse using different concentrations of picolyl azide in the click reaction, with the selected concentration of 0.1 µM noted. (F) Flow cytometry data comparing EdU and DNA fluorescence intensity in the presence of 2% NCS or 4% FBS, showing no differences in staining or the proportion of EdU-labeled cells. (G) Flow cytometry analysis of EdU-labeled C2C12 cells, evaluating DNA fluorescence intensity with various DNA dyes and concentrations.

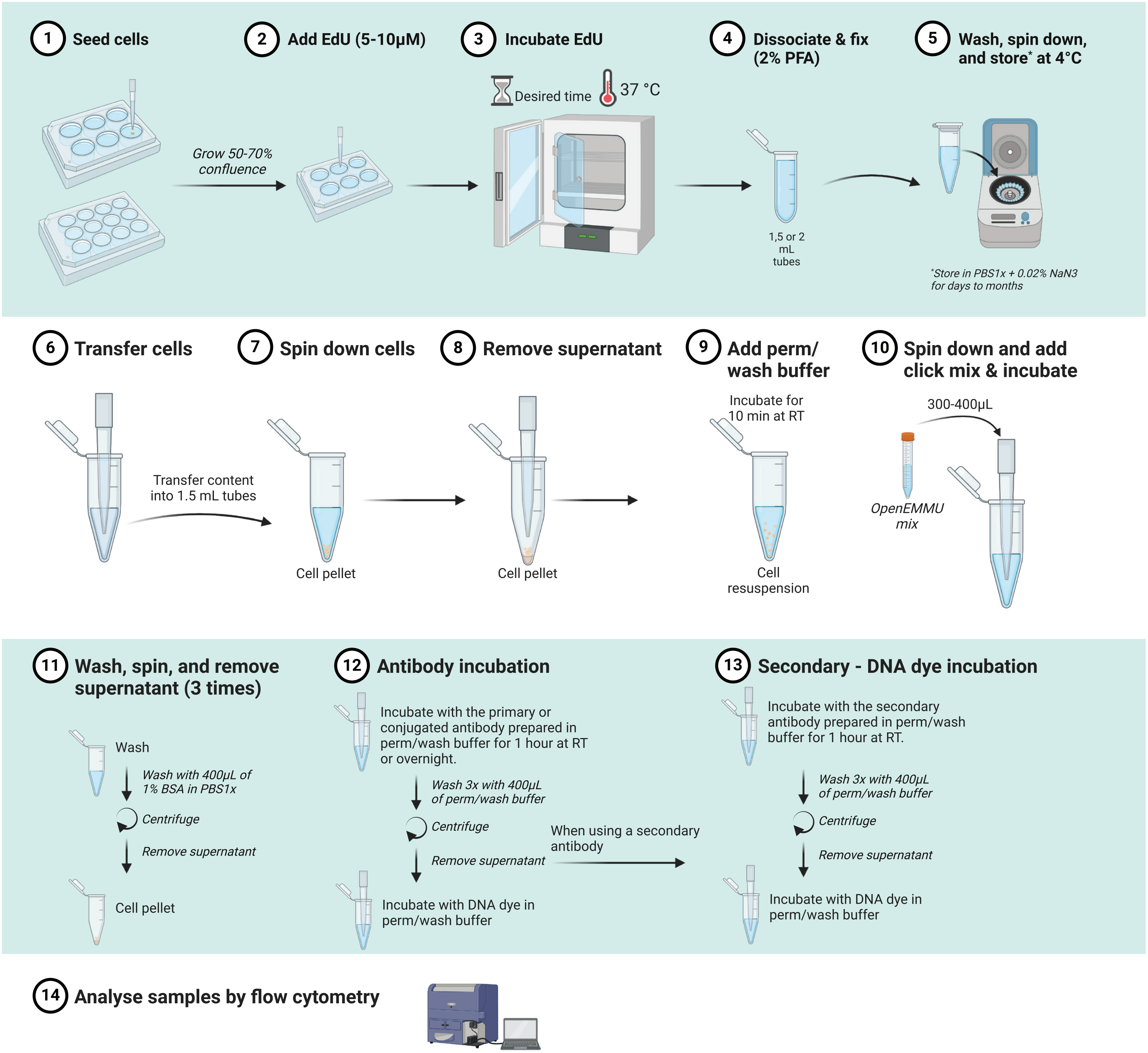


**Fig. S2. Detailed protocol for OpenEMMU click-based detection of DNA-replicating cells via flow cytometry.** Step-by-step guide for performing flow cytometry to detect DNA-replicating cells labeled with EdU using OpenEMMU. The protocol outlines 14 defined steps, designed to simplify implementation and ensure reproducibility in diverse research laboratory settings.

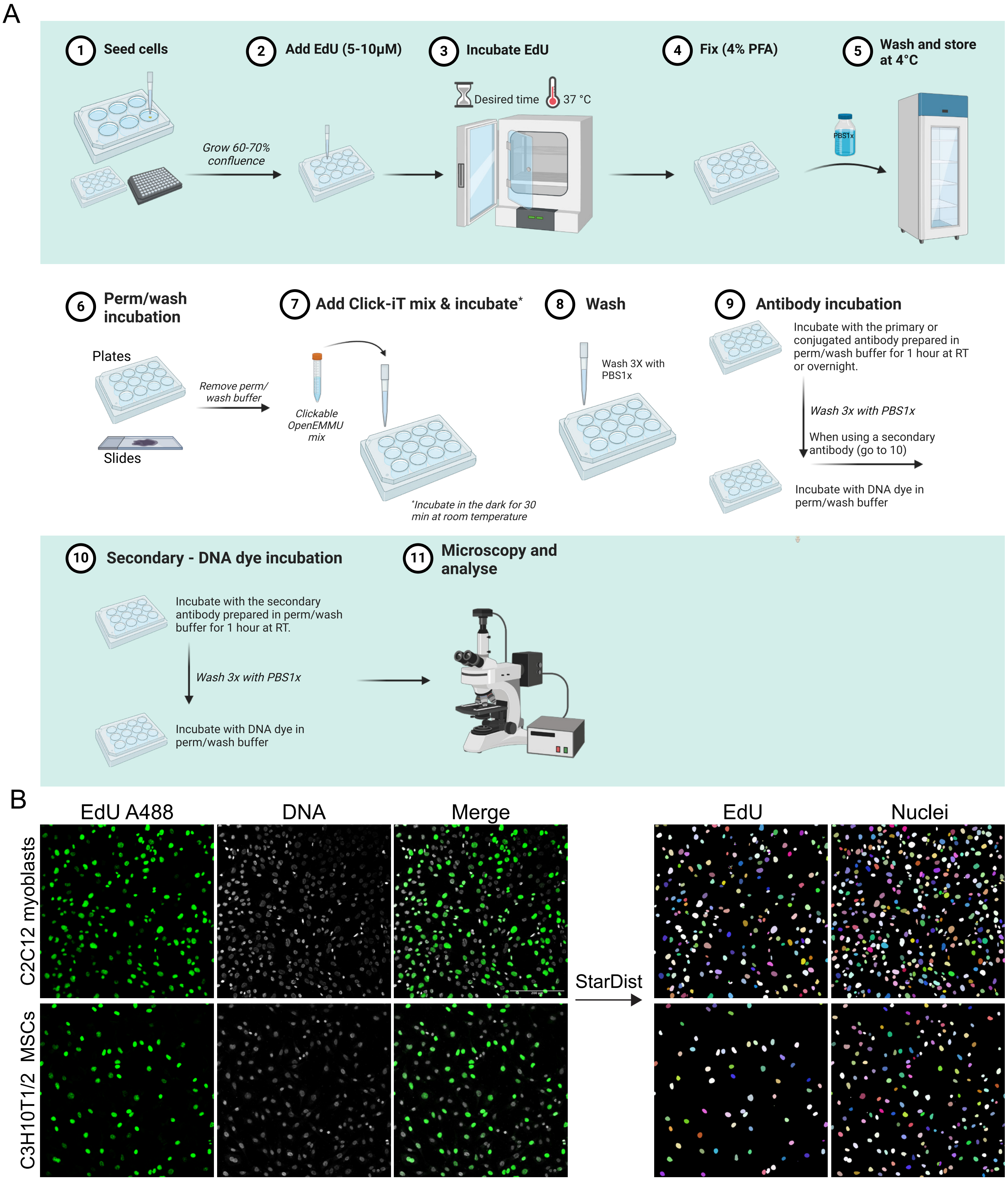


**Fig. S3. Detailed protocol for OpenEMMU click-based detection of DNA-replicating cells via imaging in cell culture plates and slides.** (A) Step-by-step protocol for imaging DNA-replicating cells labeled with EdU using OpenEMMU. The guide provides 11 defined steps to facilitate straightforward implementation and ensure reproducibility across research laboratories. (B) Confocal imaging of EdU-labelled C2C12 myoblasts and C3H10T1/2 mesenchymal cells following a 2-hour EdU uptake in growth media. Nuclei were also stained, and segmentation was performed using the StarDist plugin in Fiji, showcasing OpenEMMU's applicability in high-throughput imaging workflows for analyzing DNA replication and cell proliferation.

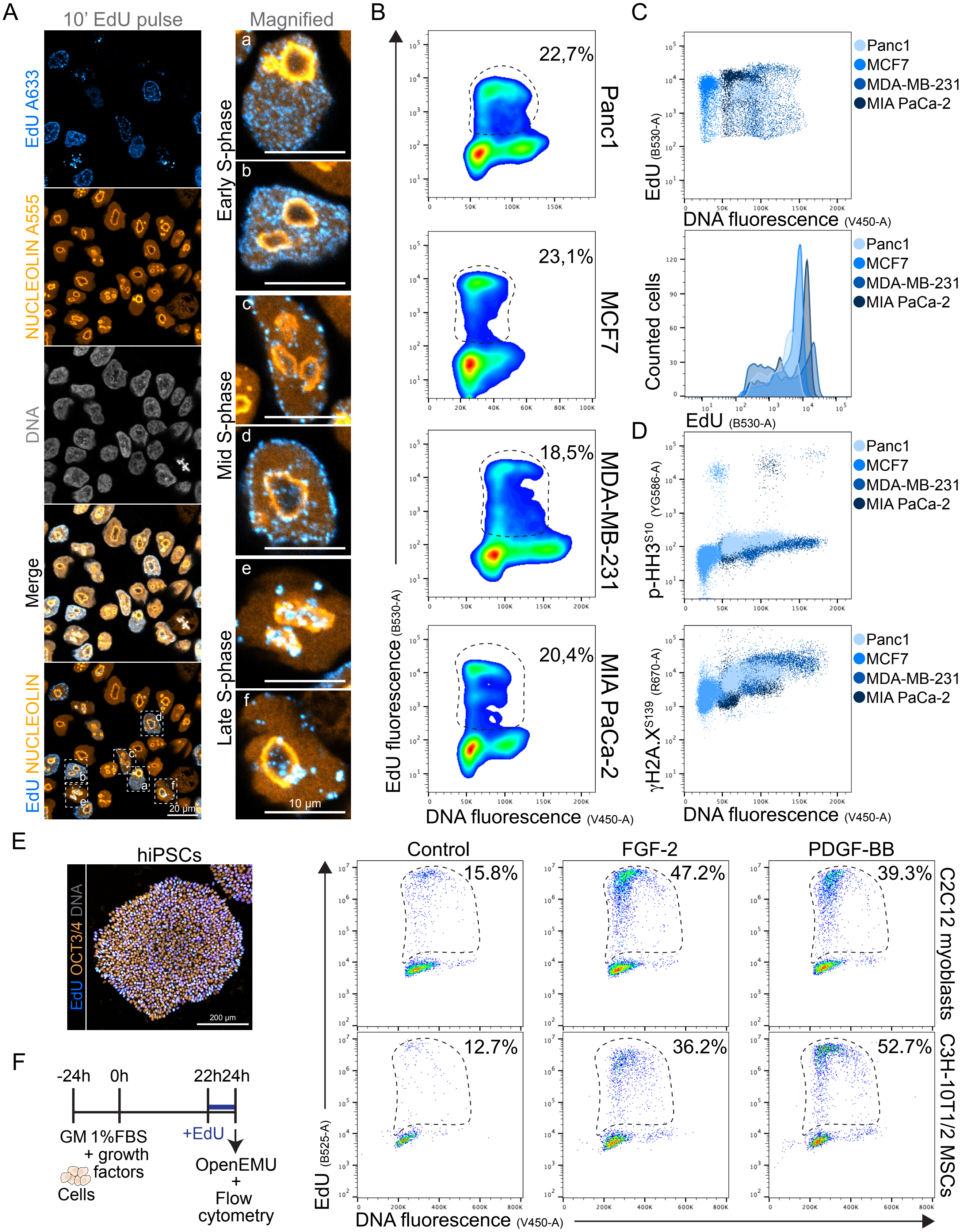


**Fig. S4. Application of OpenEMMU to normal and cancer cell lines for analysis of DNA replication.** (A) Confocal microscopy images of EdU+ hiPSCs after a 10-minute EdU pulse, with NUCLEOLIN-mediated nucleolar immunolabeling and DNA staining. The right panels show magnified cells at different S-phase stages. (B) Flow cytometry results show the proportion of EdU-labeled cancer cells after a 2-hour pulse. (C) Relationship between EdU fluorescence intensity and DNA content of different cancer cell types. (D) Flow cytometry results showing p-HH3$^{S10}$ and γH2A.X$^{S139}$ labeling after a 2-hour EdU pulse in which EdU was also co-stained. (E) Confocal microscopy images showing a high proportion of hiPSCs in S-phase after a 2-hour EdU pulse, with OCT3/4 immunolabeling and DNA staining. (F) Flow cytometry analysis of EdU labelling combined with DNA fluorescence in control cells and cells treated with PDGF-BB or FGF-2 for 24 hours.

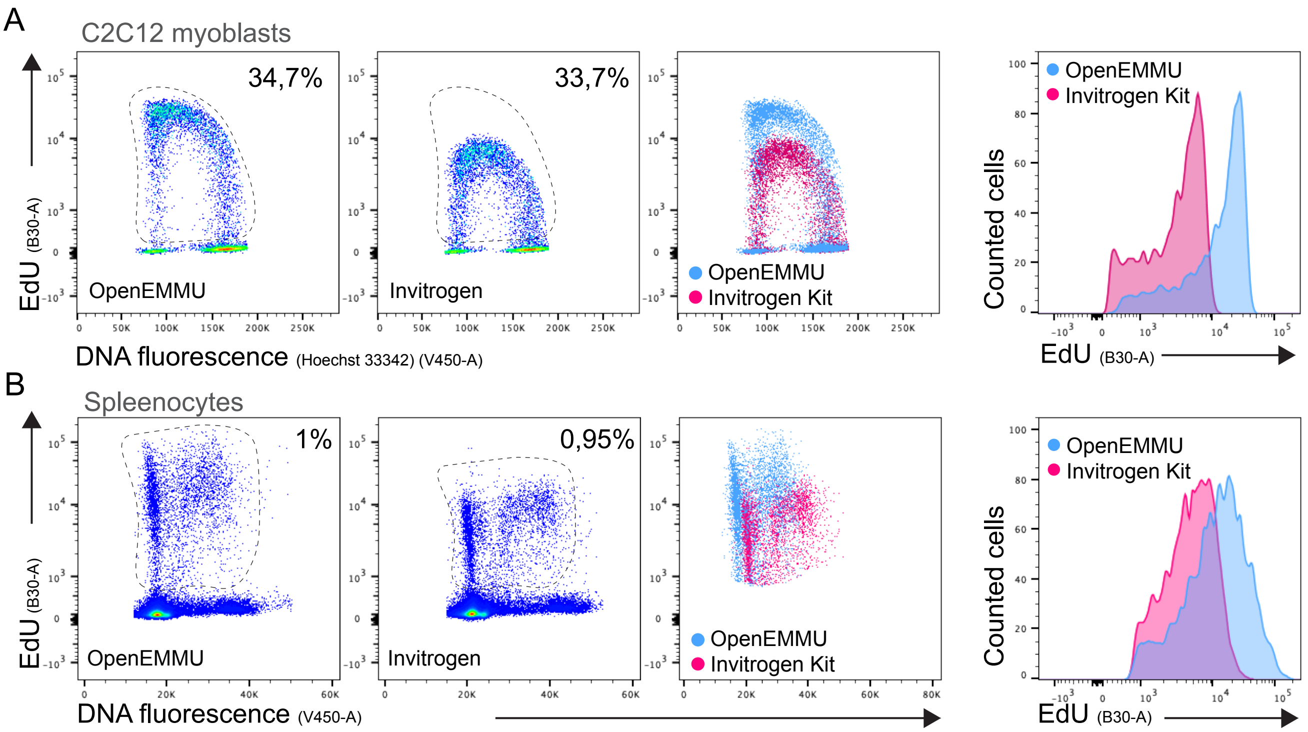


**Fig. S5. OpenEMMU outperforms commercial kits in evaluating DNA replication and S-phase of the cell cycle.** (A) Flow cytometry results comparing OpenEMMU and the Invitrogen kit in terms of EdU fluorescence intensity and the proportion of EdU-labeled cells after a 2-hour pulse in C2C12 myoblasts. (B) Flow cytometry results comparing EdU fluorescence intensity and the proportion of EdU+ cells in spleenocytes.

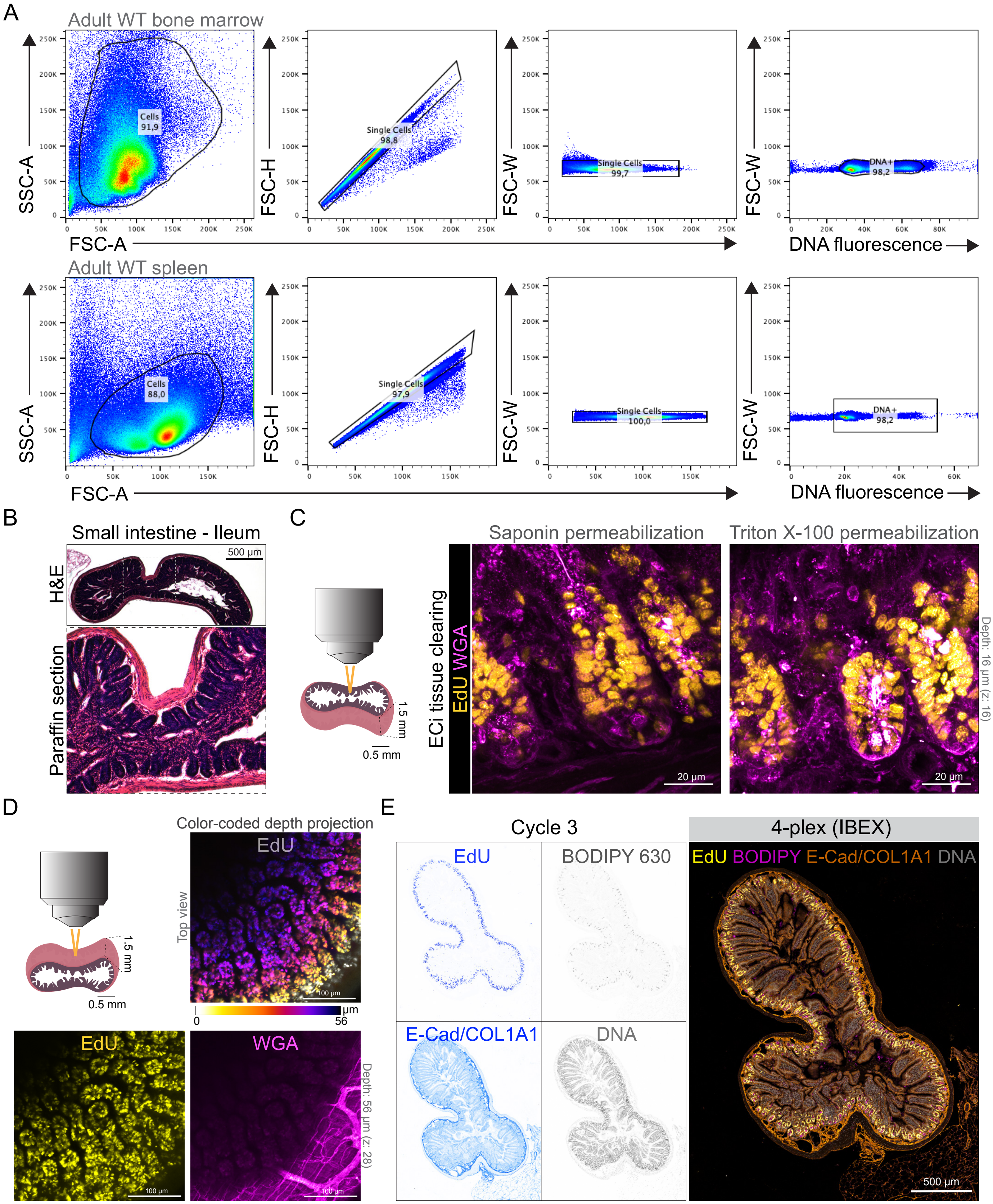


**Fig. S6. OpenEMMU for measuring DNA replication in adult organs and tissues in mice.** (A) Gating strategy employed to detect DNA replication in isolated bone marrow cells (top panels) and spleenocytes (bottom panels). The lines indicate the selected gate. (B) Hematoxylin and eosin staining of an ileum paraffin section of the small intestine. (C) OpenEMMU is compatible with ethyl cinnamate (ECi) tissue clearing and also saponin or Triton X-100 permeabilization and 3D confocal laser imaging. EdU and WGA stainings are shown. Several highly proliferating cell regions are shown, including the adult stem cell crypt and the transient amplifying cells in the villi. (D) Z-stack confocal laser imaging of EdU signals (color-coded depth projection) in DNA-replicating cells, along with WGA labeling to highlight the smooth-muscle cell layer, which is rich in blood vessels. (E) Multiplexed imaging (cycle 3) of the small intestine related to Fig. 4C.

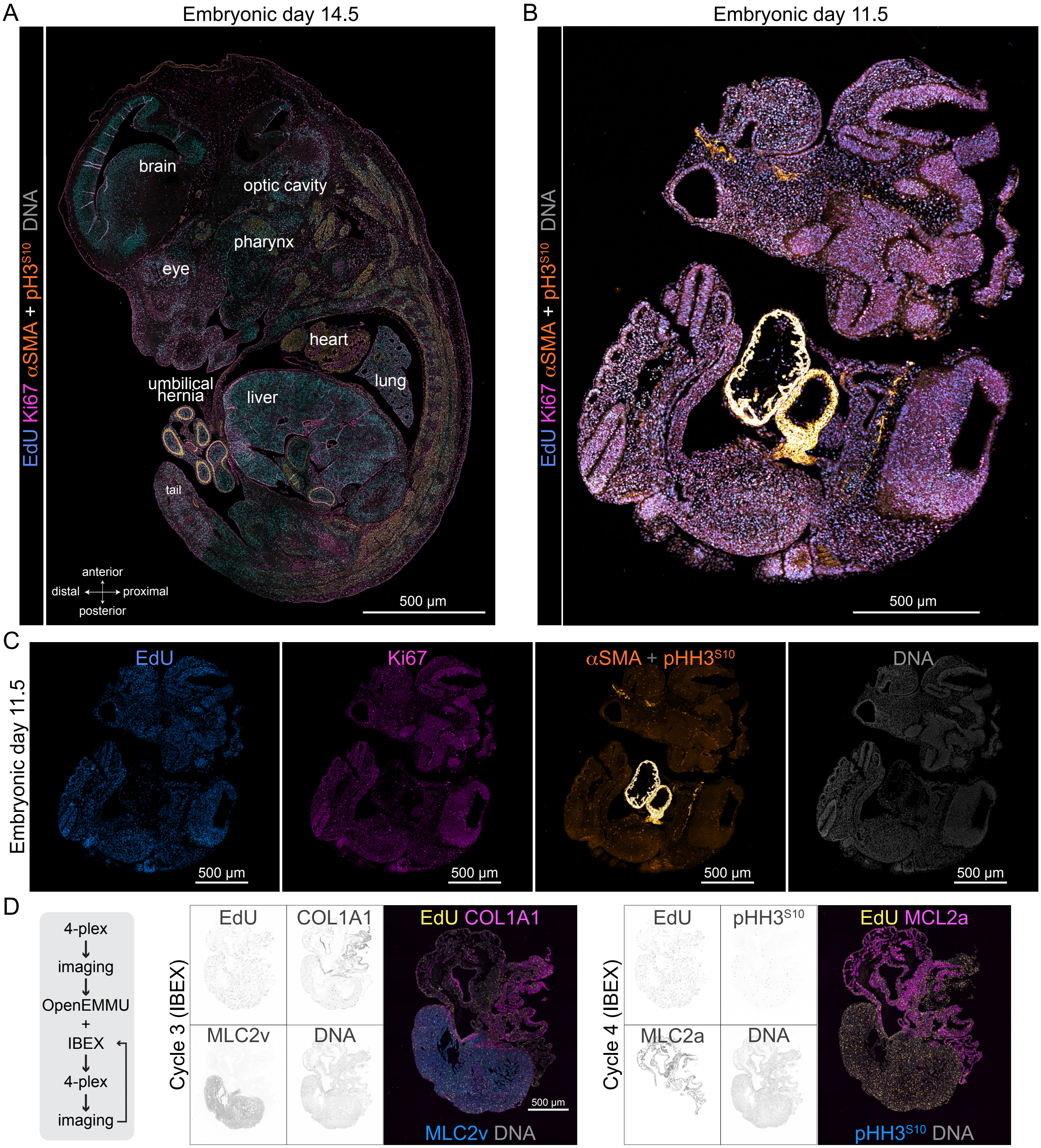


**Fig. S7. Whole-embryo and organ-specific analysis of DNA replication and mitosis in developing mouse embryos and organs.** (A) FFPE-processed embryonic day 14 mouse embryo and imaged with Thunder tile imaging of EdU signal in DNA replicating cells multiplexed with Ki67, αSMA-Cy3, pHH3$^{S10}$-PE, and DNA labeling. αSMA immunolabelling highlights the developing musculature, heart cardiomyocytes, and smooth muscle cells across multiple organs and tissues. (C) FFPE-processed embryonic day 11.5 mouse embryo and imaged with confocal tile imaging of EdU signal in DNA replicating cells, multiplexed with Ki67, αSMA, and DNA labeling. (C) Individual channels of each staining are shown in panel B. (D) FFPE section of an E14.5 heart, visualized using tiling confocal microscopy. The section underwent four rounds of immunostaining (4-plex, cycle 3; and 4-plex, cycle 4), with OpenEMMU applied during the second cycle, followed by IBEX starting from the second cycle onwards.

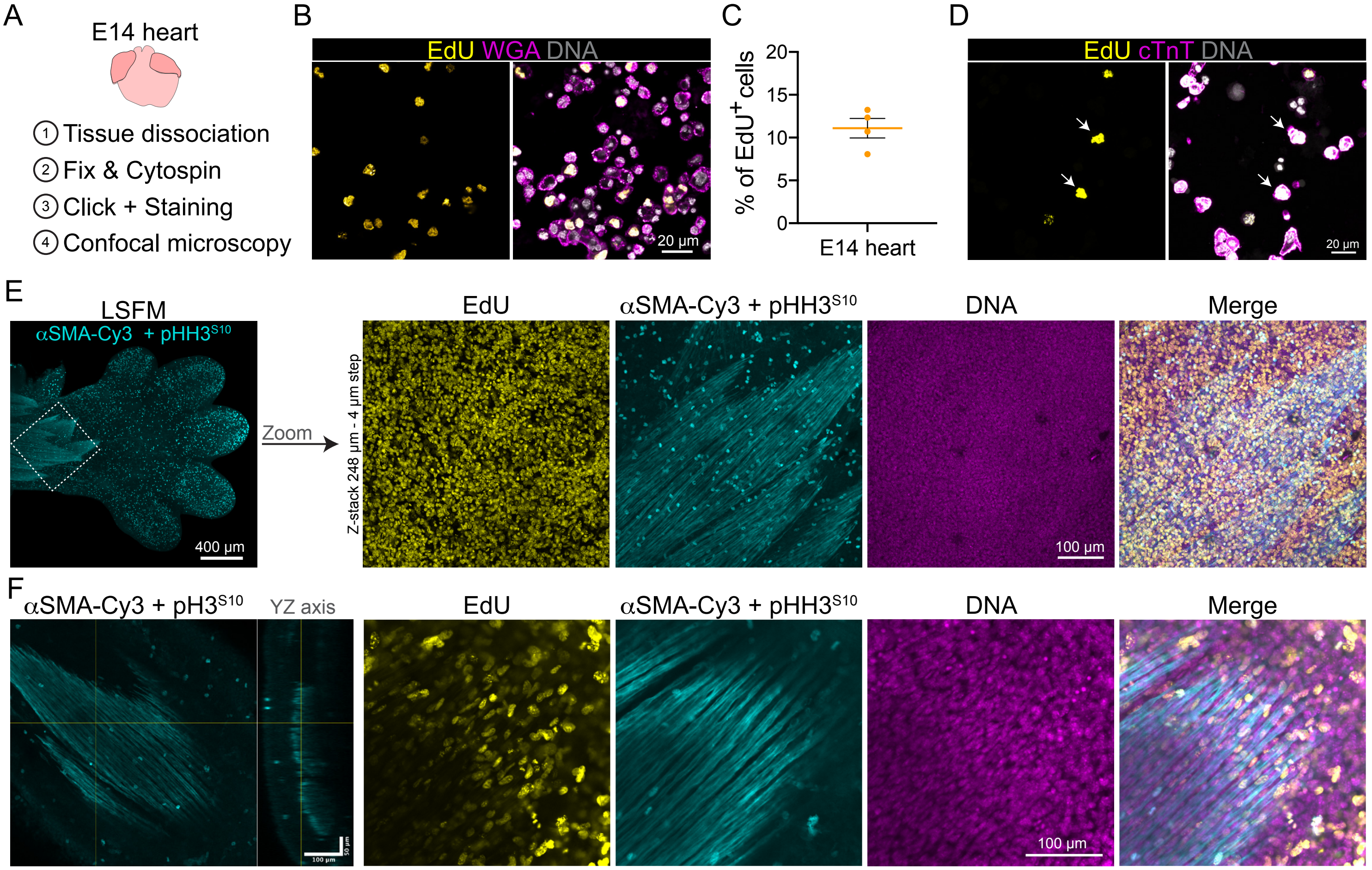


**Fig. S8. Application of OpenEMMU to embryonic heart cells and 3D imaging of forelimb development.** (A) Diagram of an embryonic day 14 (E14) mouse heart, illustrating the steps involved in the OpenEMMU protocol for single-cell imaging. (B) Confocal microscopy images of EdU+ cardiac cells following a 3-hour EdU pulse, with WGA labeling and DNA staining. (C) Proportion of EdU-labelled cells in an E14 mouse heart (n=4). (D) Confocal microscopy images of EdU-labelled cardiomyocytes (cTnT+) after a 3-hour EdU pulse, with DNA staining. The arrows show a couple of EdU+ cardiomyocytes. (E) Light sheet fluorescence imaging of EdU multiplexed with αSMA-Cy3 and pHH3$^{S10}$-PE immunolabelling, along with DNA staining of an E14 forelimb, shown from a dorsal view (same as Fig. 6D). The right panels show magnified images of each separated staining, demonstrating the high resolution of EdU labeling in DNA-replicating cells as well as immunolabelling. (F) YZ-axis light sheet fluorescence imaging showing that tissue clearing combined with OpenEMMU enables deep imaging, allowing visualization of extensor myofibers, DNA-replicating cells, and mitotic cells.

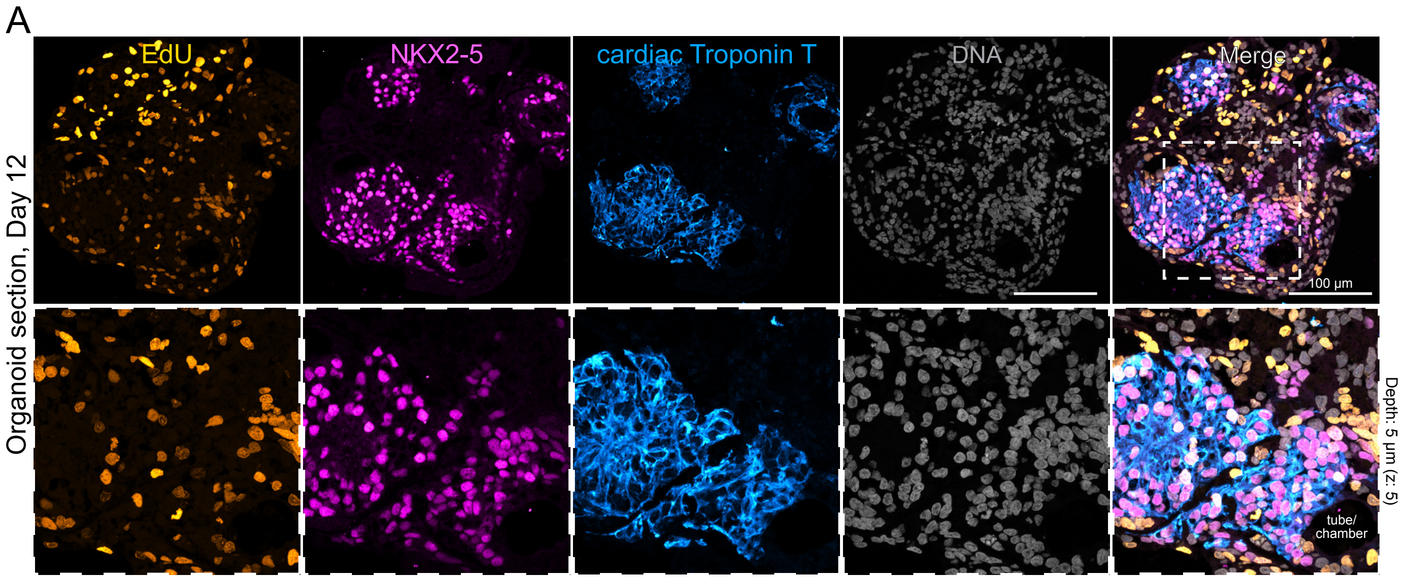


**Fig. S9. DNA replication in self-organizing 3D human cardiac organoids.** (A) FFPE-processed day 12 hCO and confocal tile imaging of EdU fluorescence in DNA replicating cells. NKX2-5 and cTnT expressing cardiomyocytes are highlighted in the boxed area. NKX2-5/EdU double-positive cells are also shown (magnified box). A tube-like cardiac chamber is also shown (bottom panels).

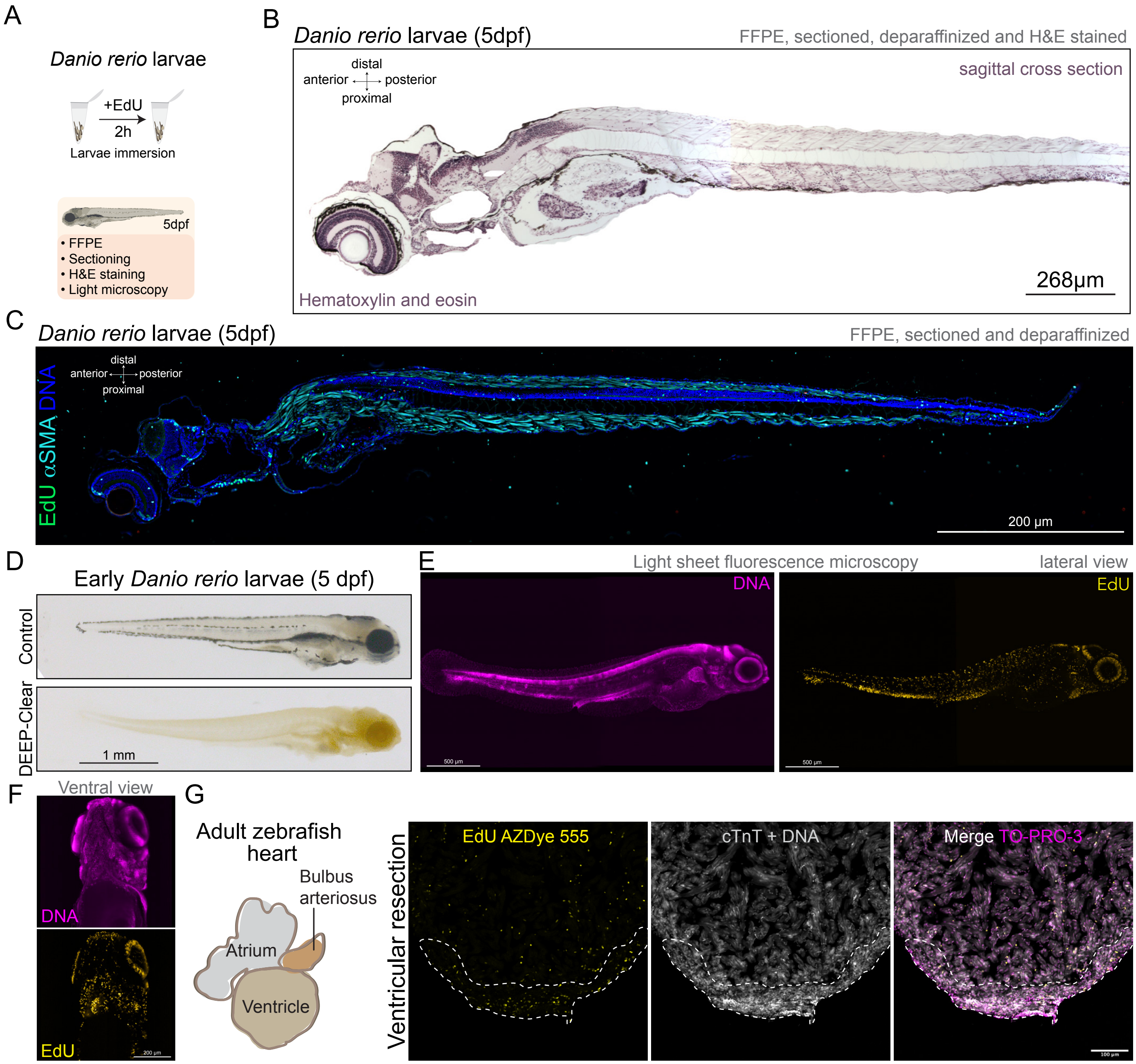


**Fig. S10. DNA replication in zebrafish larvae and under apical transection of an adult zebrafish heart.** (A) Immersion protocol aimed at labeling DNA synthesis with EdU for 2h in 5dpf developing zebrafish larvae, and processing using FFPE. (B) FFPE-processed larva sagittal cross section and light imaging of H&E staining. (C) FFPE-processed larva sagittal cross section (same as Fig. 8B) and tile imaging with Thunder Leica computational clearing of EdU signal in DNA replicating cells. αSMA immunolabeling depicts the developing musculature and growing myotomes. Nuclei were stained for DNA. (D) Whole-mount micrographs showing a growing *Danio rerio* 5dpf larva with or without DEEP-Clearing. (E, F) LSFM of a whole 5dpf zebrafish larva labeled with EdU and DNA dye, showing a lateral (E) and ventral view (F). (G) EdU, cTnT-AF647, and DNA labeling revealing proliferative cells after 7 days of apical ventricular resection in an adult zebrafish heart, visualized by confocal microscopy. The dashed lines surround the regenerating apical area of the ventricle.
