## Supplementary material for "OpenEMMU: a versatile, open-source EdU multiplexing methodology for studying DNA replication and cell cycle dynamics": Table S1

**Supplementary Materials**

**This PDF file includes:**

**Figs. S1 to S10**

**Legends for tables S1**

**Table S1. Key resources table.**

| **Reagent type (species) or resource** | **Designation** | **Source or reference** | **Identifiers** | **Additional information** |
| --- | --- | --- | --- | --- |
| Antibody | Alexa Fluor® 647 anti-Tubulin-α Antibody | BioLegend | AB_2563178 (Cat. No. 627908) | Immunofluorescence (1:250) |
| Antibody | PE anti-Histone H3 Phospho (Ser10) Antibody | BioLegend | AB_2564562 (Cat. No. 650807) | Immunofluorescence (1:250) |
| Antibody | Nucleolin (D4C7O) Rabbit mAb #14574 | Cell Signaling Technology | (Cat. No. 14574) | Immunofluorescence (1:500) |
| Antibody | Oct-3/4 (H-134) | Santa Cruz Biotechnology | sc-9081 | Immunofluorescence (1:250) |
| Antibody | BD Horizon™  Purified Rat Anti-Mouse CD16/CD32 antibody | BD Bioscience | Mouse BD Fc Block™ (Cat. No. 553141) | Flow cytometry (1:100) |
| Antibody | BD Horizon™ BV786 Rat Anti-Mouse CD45 (30-F11) | BD Bioscience | AB_2716861 (Cat. No. 564225) | Flow cytometry (1:200) |
| Antibody | BD Pharmingen™ APC-Cy™7 Rat Anti-CD11b | BD Bioscience | AB_396772 (Cat. No. 561039) | Flow cytometry (1:250) |
| Antibody | BD Horizon™ BB700 Hamster Anti-Mouse CD11C | BD Bioscience | AB_2869773 (Cat. No. 566505) | Flow cytometry (1:250) |
| Antibody | BD Horizon™ BV786 Mouse Anti-Human CD3 | BD Bioscience | AB_2869863 (Cat. No. 566781) | Flow cytometry (1:50) |
| Antibody | BD Horizon™ BUV395 Mouse Anti-Human CD4 | BD Bioscience | AB_2738273 (Cat. No. 563552) | Flow cytometry (1:50) |
| Antibody | BD Horizon™ BUV805 Mouse Anti-Human CD8 | BD Bioscience | AB_2833078 (Cat. No. 612889) | Flow cytometry (1:50) |
| Antibody | BD™ APC Mouse Anti-Human CD25 | BD Bioscience | AB_400551 (Cat. No. 340939) | Flow cytometry (1:50) |
| Antibody | Anti-Ki67 antibody (ab15580) | Abcam | Rabbit Polyclonal (Cat. No. ab15580) | Immunofluorescence (1:250), 1:100 for 3D |
| Antibody | Cardiac Troponin T (cTnT) Mouse Monoclonal Antibody (13-11) | ThermoFisher Scientific | AB_11000742 (Cat. No.  MA5-12960) | Immunofluorescence (1:500),  1:250 for 3D |
| Antibody | Ki-67 Antibody, anti-human/mouse, Vio® R667, REAfinity™ | Miltenti Biotec | REA 183  (Cat. No. 130-120-422) | Immunofluorescence (1:100),  1:50 for 3D |
| Antibody | Anti-Actin, α-Smooth Muscle-Cy3™ antibody, Mouse monoclonal | Sigma-Aldrich | Cat. No. C6198 | Immunofluorescence (1:250) |
| Peptide | Flash Phalloidin™ Red 594 | BioLegend | Cat. No. 424203 | Staining (1:100) |
| Antibody | β-Catenin Antibody, anti-human/mouse, PE, REAfinity™ | Miltenti Biotec | REA480  (Cat. No. 130-122-006) | Immunofluorescence (1:100) |
| Antibody | BD Pharmingen™ Alexa Fluor® 647 Mouse Anti-Cardiac Troponin T (cTnT) | BD Bioscience | AB_2739341 (Cat. No. 565744) | Immunofluorescence (1:250),  1:100 for 3D |
| Antibody | BD Pharmingen™ PE Mouse Anti-Cardiac Troponin T (cTnT) | BD Bioscience | AB_2738939 (Cat. No. 564767) | Immunofluorescence (1:250) |
| Antibody | Alexa Fluor® 647 anti-human/mouse/rat PCNA Antibody, Mouse IgG2a, κ, PC10 | BioLegend | AB_2267947 (Cat. No. 307912) | Immunofluorescence (1:250) |
| Antibody | CD31 (PECAM-1) Monoclonal Antibody (390), PE-Cyanine7, eBioscience™ | ThermoFisher Scientific | AB_2716949 (Cat. No. 25-0311-82) | Flow cytometry (1:250) |
| Antibody | BD Pharmingen™ PE Rat Anti-Mouse CD140A, Clone APA5 | BD Bioscience | AB_2737787 (Cat. No. 562776) | Flow cytometry (1:200) |
| Antibody | PE anti-mouse CD140a Antibody | BioLegend | AB_1953268 (Cat. No. 135905) | Flow cytometry (1:200) |
| Antibody | Nkx-2.5 Antibody (H-114), rabbit polyclonal IgG | Santa Cruz Biotechnology | Cat. No. sc-14033 | Immunofluorescence (1:200) |
| Lipid stain | Bodipy BDP® 630/650 lipid stain | Lumiprobe | Cat. No. 1233-1mg | Immunofluorescence (250 nM) |
| Antibody | COL1A1 (E8F4L) XP® Rabbit mAb (Alexa Fluor® 647 Conjugate) | Cell Signaling Technology | Cat. No. 72827 | Immunofluorescence (1:100) |
| Antibody | E-Cadherin (24E10), Rabbit mAb | Cell Signaling Technology | Cat. No. 3195T | Immunofluorescence (1:250) |
| Antibody | MLC2v Antibody, anti-human/mouse/rat, PE, REAfinity™ | Miltenti Biotec | REA401  (Cat. No. 130-119-680) | Immunofluorescence (1:100) |
| Antibody | MLC2a Antibody, anti-human/mouse/rat, APC, REAfinity™ | Miltenti Biotec | REA398  (Cat. No. 130-118-674) | Immunofluorescence (1:100) |
| Clearing agent | N,N,N′,N′-Tetrakis(2-hydroxyethyl)ethylenediamine, THEED | Sigma-Aldrich | Cat. No. 87600 | 9% (v/v) in distilled water |
| Picolyl Azide, fluorescent probe | AZDye 488 Picolyl Azide | Click Chemistry Tools | CCT-1276 | Detailed in text and Materials and Methods |
| Picolyl Azide, fluorescent probe | AZDye 555 Picolyl Azide | Click Chemistry Tools | CCT-1288 | Detailed in text and Materials and Methods |
| Picolyl Azide, fluorescent probe | AZDye 633 Picolyl Azide | Click Chemistry Tools | CCT-1549 | Detailed in text and Materials and Methods |
| Picolyl Azide, fluorescent probe | AZDye 680 Picolyl Azide | Click Chemistry Tools | CCT-1511 | Detailed in text and Materials and Methods |
| Alexa Fluor, secondary antibody | Various | ThermoFisher Scientific |  | Immunofluorescence (1:500), 1:250 for 3D |
